# INPP4B drives lysosome biogenesis to restrict leukemic stem cell differentiation and promote leukemogenesis

**DOI:** 10.1101/2021.03.25.437029

**Authors:** John F. Woolley, Keyue Chen, Golam T. Saffi, Gizem E. Genc, Daniel K.C. Lee, Irakli Dzneladze, Ruijuan He, Jonathan T.S. Chow, Martino M. Gabra, Meong Hi Son, Ché M.P. Melo, Candaice A. Newell, Aobo He, Erwin M. Schoof, Stephanie Z. Xie, Emily M. Mangialardi, Max Kotlyar, Ayesha Rashid, Miki. S. Gams, Jean Vacher, Cynthia J. Guidos, Igor Jurisica, John E. Dick, Roberto J. Botelho, Mark D. Minden, Leonardo Salmena

**Author notes:** Department of Pharmacology & Therapeutics, Institute of Systems, Molecular and Integrative Biology, University of Liverpool, UK. Department of Pediatrics, Samsung Medical Center, Sungkyunkwan University School of Medicine, Seoul, Korea (Meong Hi Son). Correspondence (L.S.).

## Abstract

Signaling pathways that control vital features of leukemic stem cells including multipotency, self-renewal, clonal expansion and quiescence remain unclear. Emerging studies illustrate critical roles for lysosomes in hematopoietic and leukemic stem cell fate. By investigating consequences of *INPP4B* alterations in AML, we have discovered its role in driving leukemic ‘stemness’. We observed that *INPP4B* is highly expressed leukemic stem cell populations and *Inpp4b*-deficeint leukemias demonstrate increased disease latency, reduced leukemia initiating potential which is associated with a differentiated leukemic phenotype. Molecular analyses show that *Inpp4b*-deficient leukemias have compromised lysosomal gene expression, lysosomal content, and lysosomal activity. Our discovery of a novel pathway linking INPP4B, lysosomal biogenesis and leukemic stemness, provides a mechanism to explain the association of high *INPP4B* expression with poor AML prognosis, and highlights novel patient stratification strategies and LSC-specific leukemic therapies.

**Key Points:**

Our findings highlight a novel pathway linking INPP4B, lysosomal function and leukemic stemness that explains the prognostic role of INPP4B in AML.
Our data reveal the utility of INPP4B as a biomarker of aggressive AML and provide a rationale to explore INPP4B and its associated function in lysosome biology as novel strategies to target LSC and AML

## Introduction

Acute myeloid leukemia (AML) is an aggressive malignancy of the bone marrow characterized by dysregulated proliferation of immature myeloid cells and poor survival rates (Papaemmanuil et al., 2016; Siegel et al., 2018). AML exists in a cellular hierarchy with leukemic stem cells (LSC) at the pinnacle that possess ‘stemness’ attributes including a capacity for self-renewal, multipotent differentiation, long-term clonal propagation and quiescence which sustain the malignant undifferentiated myeloid ‘blast’ cells that define AML and drive relapse (Dick, 2008; Kreso and Dick, 2014). Despite an increased understanding of the roles for ‘stemness’ in relapse and chemoresistance in AML over the past decades, the underlying molecular mechanisms remain unclear.

We previously reported that expression of *Inositol Polyphosphate-4-Phosphatase Type II* (*INPP4B*), a lipid phosphatase that hydrolyzes phosphatidylinositol-3,4-bisphosphate [PtdIns(3,4)P2] to generate phosphatidylinositol-3-monophosphate [PtdIns(3)P], is elevated in the leukemic blasts of approximately one quarter of all AML patients and is associated with poor prognosis (Dzneladze et al., 2015). The notion that high INPP4B expression can drive tumorigenesis was supported by several studies studies in AML (Dzneladze et al., 2015; Recher, 2015; Rijal et al., 2015; Wang et al., 2016; Zhang et al., 2017) and other malignancies (Chi et al., 2015; Gasser et al., 2014; Guo et al., 2016; Rodgers et al., 2021). These findings prompted us to investigate specific tumour-promoting roles for INPP4B in leukemogenesis. Herein we present lines of evidence illustrating a novel role for INPP4B in the regulation of lysosomal properties, through which it controls leukemia stem cell maintenance and AML phenotypes.

## Results

### High levels *INPP4B* expression in AML are associated with leukemic stemness signatures

Analysis of overall (OS) and event-free survival (EFS) demonstrate that *INPP4B* transcript levels effectively stratify AML patients (**Fig. 1A, B)** as previously demonstrated (Dzneladze et al., 2015, 2018; Rijal et al., 2015). We identified the most significantly overrepresented gene ontologies associated with the top 200 genes enriched in high *INPP4B* patient samples across several AML patient databases (**Supplementary Table 1)**. This analysis highlighted significant associations with phosphoinositide, PI3K and Akt signal transduction and notably, signaling networks related to leukemic stemness including hemopoiesis, stem cell proliferation, megakaryocyte differentiation, and hematopoietic progenitor cell differentiation (**Fig. 1C).** Given these associations, we specifically explored the association of *INPP4B* expression with leukemic stemness by measuring specific enrichment of the LSC-17 signature (Ng et al., 2016) where we observed a striking enrichment in patient samples with high INPP4B expression across all AML patient databases tested (**Fig. 1D, E & Supplementary Fig. 1 A, B**) (Balgobind et al., 2011; Cancer Genome Atlas Research Network et al., 2013; Klein et al., 2009; Metzeler et al., 2008; Verhaak et al., 2009). Assessment of *INPP4B* in gene expression profiles from 227 AML patients’ blast cells sorted based on the expression of CD34 and CD38 surface markers demonstrated highest levels in CD34^+^CD38^-^ populations and lower expression within other subpopulations (**Fig. 1F**. Additionally, cell populations defined functionally through bone marrow transplant experiments (Eppert et al., 2011; Ng et al., 2016) revealed that leukemic populations with disease initiating capacity (LSC^+^) had higher levels of *INPP4B* transcript compared to leukemic lineages that did not generate disease (LSC^-^) (**Fig. 1G**). These data represent the first evidence pointing to an association between *INPP4B* and leukemic stemness.

**Figure 1.**
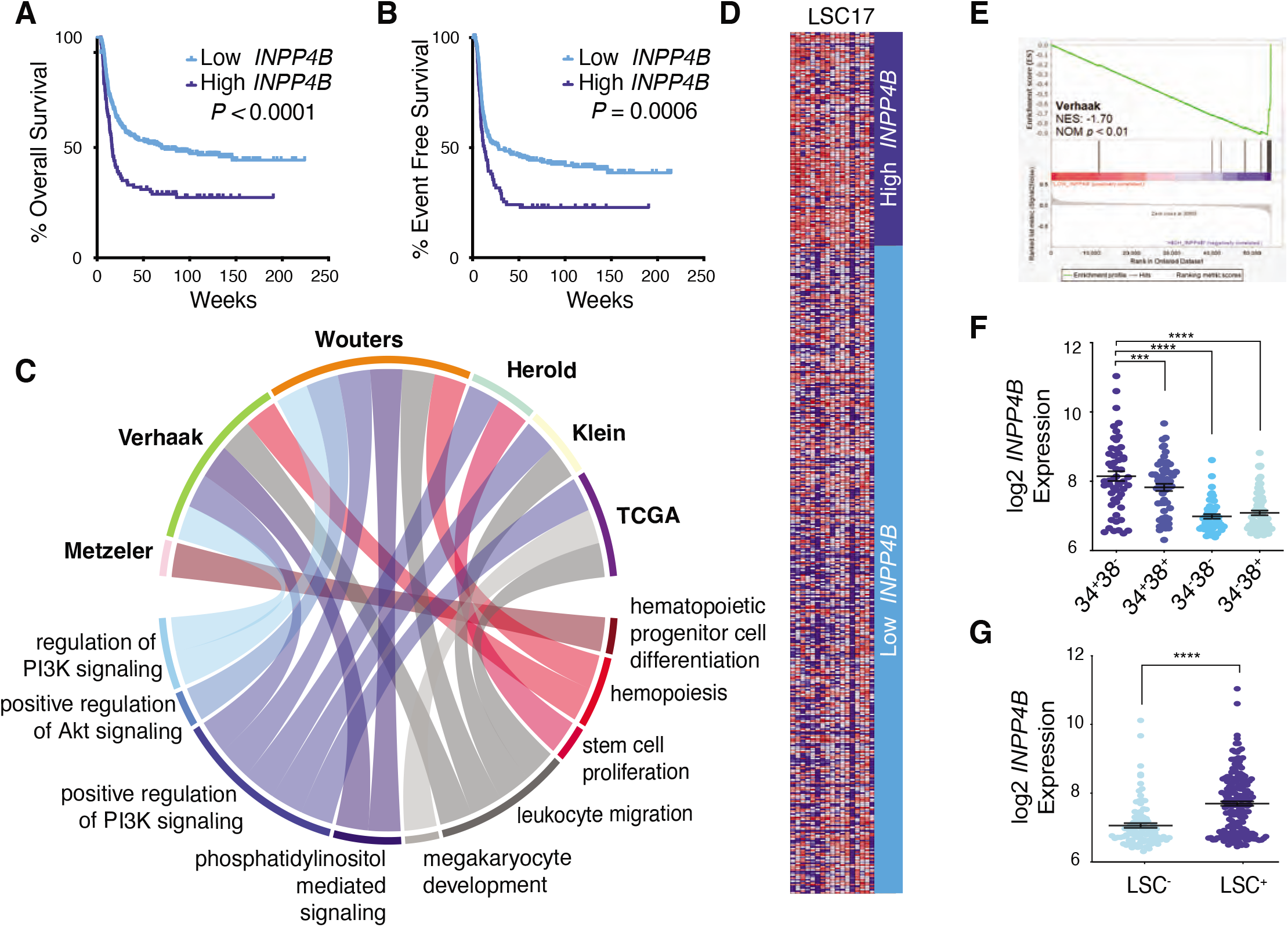
High levels INPP4B expression are associated with stemness signatures in leukemia and hematopoietic progenitor cells. A. Kaplan Meier analysis of overall survival in INPP4B^high^ vs. INPP4B^low^ AML patients from Verhaak dataset. B. Kaplan Meier analysis of event free survival in INPP4B^high^ vs. INPP4B^low^ AML patients from Verhaak dataset. C. Enriched terms related to phosphatidylinositol signaling or the hematopoietic compartment stem cells were highlighted in a chord diagram that was generated with the circlize R package. D. Heatmaps of LSC-17 gene expression in the Verhaak AML patient dataset, ranked by INPP4B expression (INPP4B-high quartile highlighted dark blue). E. Gene set enrichment analysis (GSEA) enrichment plot of the LSC-17 gene set in the Verhaak AML patient dataset ranked by INPP4B AML expression. F. Expression of INPP4B from 227 AML patient-derived samples sorted based on the expression of combinations of CD34 and CD38 surface markers. G. INPP4B expression in 278 sorted fractions characterized as LSC+ and LSC-enriched by their engraftment potential into NOD/SCID mice.

### *Inpp4b-deficiency* in *MLLAF9* leukemias leads to increased disease latency and reduced leukemic initiating capacity

To investigate a role for *Inpp4b* in leukemogenesis *in vivo*, we generated a model of *Inpp4b*-deficient leukemia using the constitutive *Inpp4b-knockout* mouse model (*Inpp4b^-/-^*) (Ferron et al., 2011). Leukemias were generated by transducing respective LSK cells with *MSCV-MLL-AF9-Venus* retrovirus followed by tail vein injection into sub-lethally irradiated syngeneic host C57BL/6 mice **(Supplementary Fig. 2A**). Notably, all host animals transplanted with *Inpp4b^+/+^* LSK cells succumbed to leukemia associated disease, however ~40% of *Inpp4b^-/-^* LSK-transplanted mice survived beyond 300 days indicating reduced leukemia initiating potential (**Fig. 2A**). Secondary transplants demonstrated a significant increase in disease latency for the *Inpp4b^-/-^*-*MLL-AF9* leukemia when compared to the *Inpp4b^+/+^-MLL-AF9* leukemia (median survival of 76 *vs*. 41 days respectively; *P* = 0.0246 **(Fig. 2B**)). Limiting dilution cell transplantation assays (LDA) revealed that *Inpp4b^-/-^* leukemias have a significant decrease (1/147 *vs* 1/300 *P* = 0.0331) in leukemic initiating cell (LIC) frequency (**Fig. 2C)**. *Inpp4b^+/+^* and *Inpp4b^-/-^* leukemia bearing mice treated with a 7 day regimen of 100mg/kg AraC demonstrated that *Inpp4b^-/-^* leukemias are sensitized to chemotherapy. These data indicate that *Inpp4b*-deficiency in a *MLL-AF9* leukemia model generates leukemias with decreased leukemia initiating potential, increased disease latency and increased chemosensitivity.

**Figure 2.**
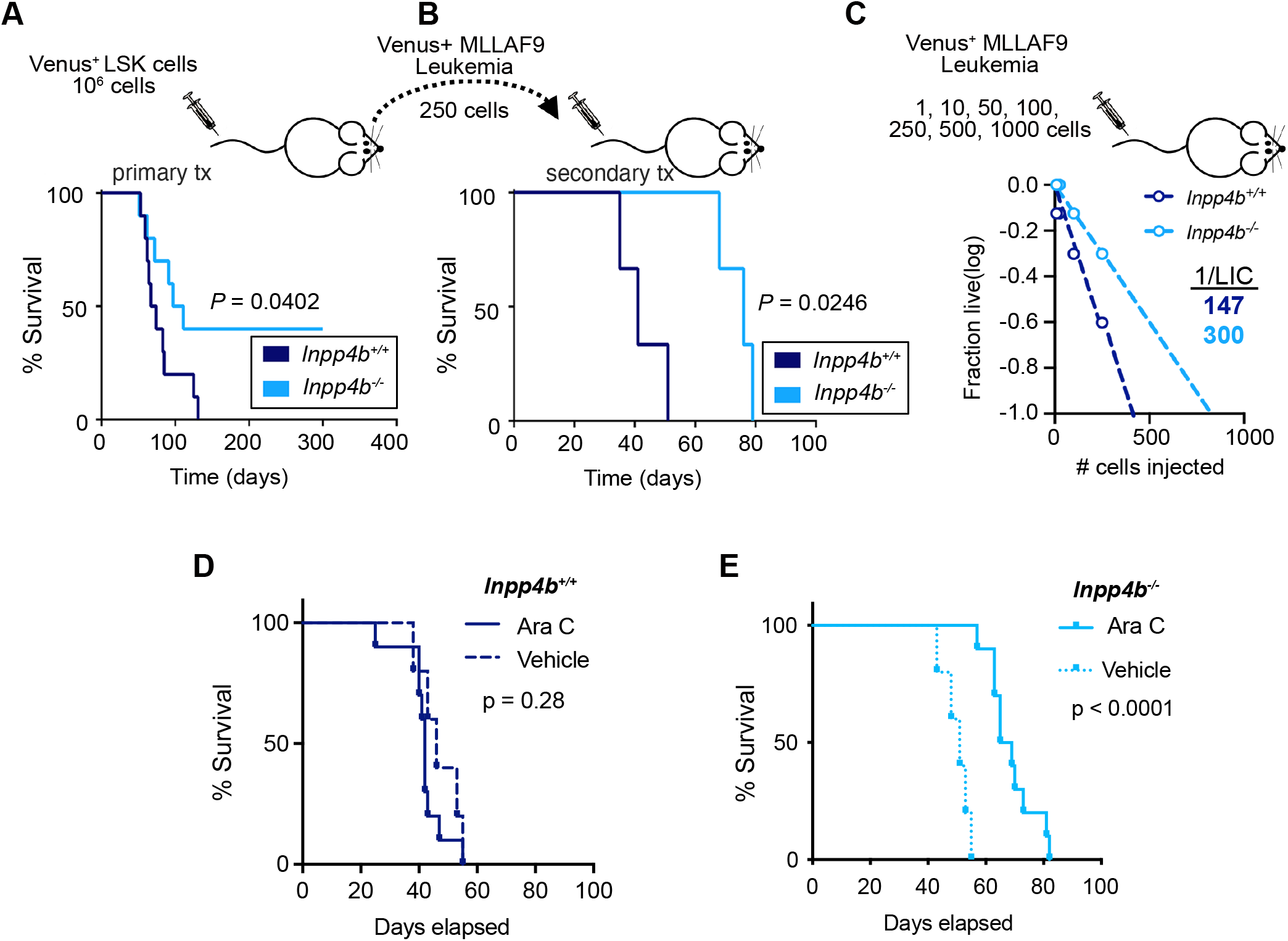
Inpp4b-deficient MLL-AF9 cells have a decreased leukemogenic potential. A. Kaplan-meier survival analysis of primary Venus+-MLL-AF9 leukemias generated from Inpp4b+/+ and Inpp4b-/- mouse LSK cells (n=10). B. Kaplan-meier survival analysis of secondary MLL-AF9 leukemias generated by transplantation of cells from Inpp4b+/+ and Inpp4b-/- MLL-AF9 leukemia blasts (n=6). C. Limiting dilution assay (LDA) of Inpp4b+/+ (dark blue (n=12)) and Inpp4b-/- MLL-AF9 (light blue (n=12)) leukemia cells (1000, 250, 100, and 25 cells) into host wildtype C57BL/6 mice. D,E. Inpp4b+/+ and Inpp4b-/- MLL-AF9 leukemia bearing mice were treated with 100mg/kg AraC daily for 7 days starting at day 14 post transplant.

### *Inpp4b*-deficiency in *MLLAF9* leukemias leads to leukemia with altered differentiation status

Leukemia forming cell (LFC) assays demonstrated that *Inpp4b*-deficiency leads to a significant decrease in total leukemic colonies (**Fig. 3A)**, characterized by a decrease in type-I colonies (spherical with defined border, resembling primitive hematopoietic colony formation) and an increase in more differentiated type-II colonies (diffuse, lacking a defined border) (**Fig. 3B,C)** (Somervaille and Cleary, 2006). Wright–Giemsa staining revealed an elevated proportion of multi-lobed, horseshoe-shaped, or diffuse nuclei, characteristic of a more differentiated blast cell phenotype in *Inpp4b^-/-^*-leukemias (**Fig. 3D**). We used mass cytometry to quantitate 15 cell-surface hematopoietic differentiation markers on *Inpp4b^-/-^* and *Inpp4b^+/+^ MLL-AF9* leukemias. Compared to *wild-type* mouse bone marrow, mice burdened with leukemia had very few erythroid and T/B lymphocytes (**Fig. Supplementary 3A**) and consisted predominantly of leukemic cells with CD11b^+^ CD16/32^+^ Gr1^-^ (**Fig. 3E,G)** cells that lacked Sca-1 and CD150 **(Fig. Supplementary 3B**), indicative of disease with myelomonocytic differentiation. Notably, a *Inpp4b^-/-^* leukemic cells expressed significantly more Gr1 compared to Inpp*4b^+/+^* leukemias (**Fig. 3E,F)**, consistent with more granulocytic differentiation. *Inpp4b^-/-^* leukemias also displayed more CD16/32^+^ CD117^+^ cells (**Fig. 3G,H**), but this difference was not significant. *Inpp4b^-/-^* and Inpp*4b^+/+^* leukemias also expressed significantly different levels of CD24, CD44 and CD16/32 (**Fig. 3I,J**), together indicating that *Inpp4b*-deficiency alters the differentiation status of *MLL-AF9* leukemias.

**Figure 3.**
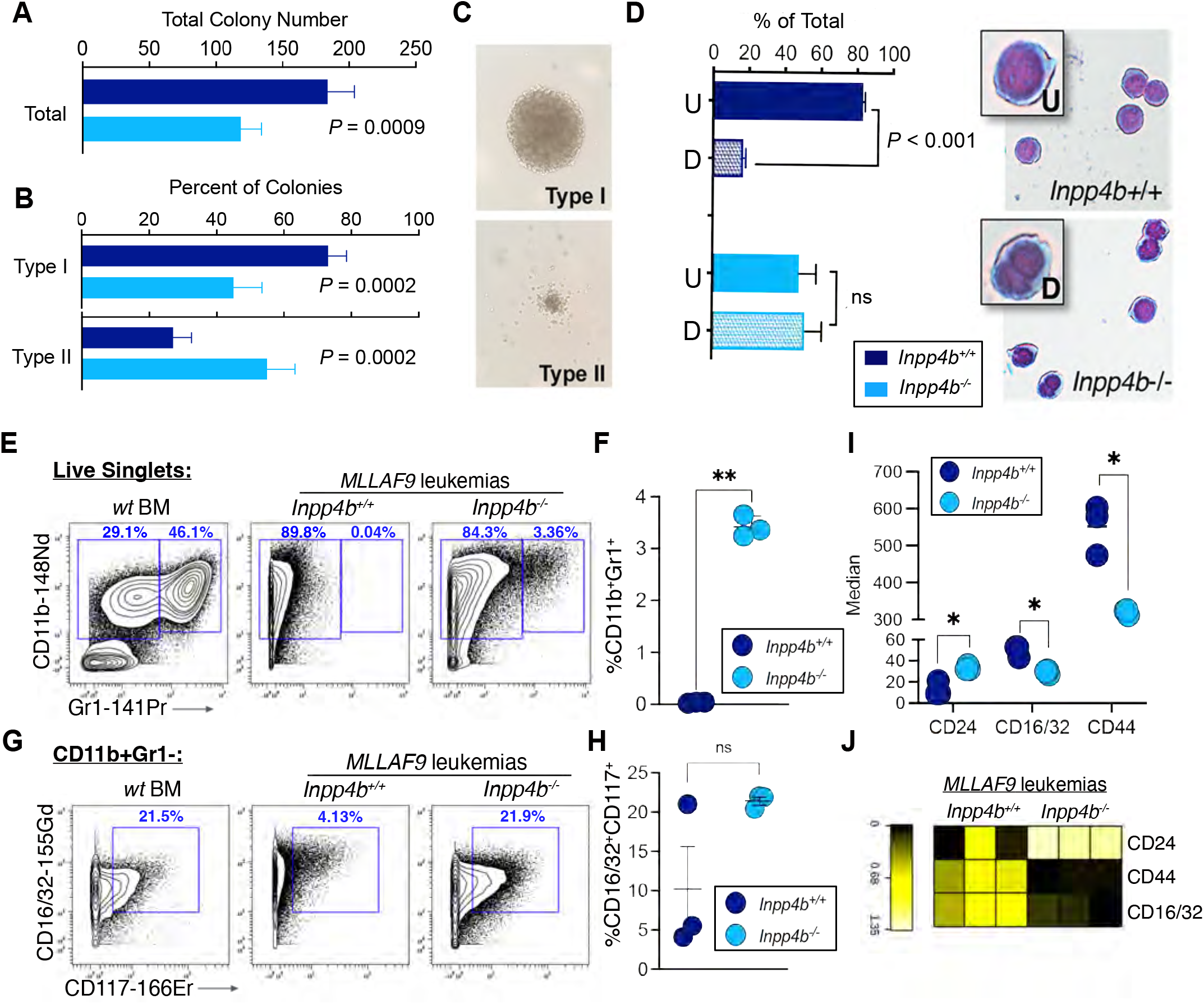
Inpp4b-deficient MLL-AF9 cells have a more differentiated phenotype. A. Total and percentage colony counts in LFC assays from Inpp4b+/+ and Inpp4B-/- murine MLL-AF9 leukemia blasts (n=8, ±S.E.M.). B. Proportion of undifferentiated and differentiated leukemic blasts from Inpp4b+/+ (n=8, ±S.E.M.). C. Representative image of type-I and type-II colonies from secondary MLL-AF9 leukemia cells. D. Representative images of Wright–Giemsa staining of leukemic blasts from Inpp4b+/+ and Inpp4b-/- MLL-AF9 terminal leukemias; (U) undifferentiated and (D) differentiated (n=3, ±S.E.M.; left). E,F. Contour plots (10% probability) show CD11b versus Gr1 expression on live singlets from Inpp4b+/+ versus Inpp4b-/- MLL-AF9 leukemia samples compared to WT BM cells (representative of n=3/group). Scatter graph shows the % CD11b+ Gr1+ cells in each AML type. Two-tailed t-test with Welch’s correction, P = 0.001. G,H. Contour plots show CD16/32 versus CD117 expression on CD11b+ Gr1-cells from representative Inpp4b+/+ and Inpp4b-/- MLL-AF9 leukemia samples. Scatter graph shows the % CD16/32+ CD117+ cells in each type of AML. Two-tailed t-test with Welch’s correction, P = 0.1726. I. Multiple unpaired t-tests with Welch’s correction. Multiple comparisons corrected by False Discovery Rate using the 2-stage step-up method of Benjamini, Krieger and Yekiuteli. *, Q=0.03 for each comparison shown. J. Heatmap visualization of the median metal intensity (MMI) of the indicated markers on CD11b+ Gr1-cells from each type of AML. Values for each marker were calculated as arcinsh (MMI(x)/scale_argument)-arcsinh (control /scale_argument) where x is the sample value, control is the lowest value for the marker across all samples, and scale argument=5.

### Identification of transcriptional networks perturbed by *Inpp4b* loss in leukemia

RNA sequencing (RNA-seq) on freshly isolated bone marrow from mice burdened with *Inpp4b^+/+^* or *Inpp4b^-/-^* leukemias revealed a total of 5462 differentially expressed genes (2434 downregulated, 3028 upregulated; FDR < 0.05; **Supplementary Fig. 4A, B**, **Supplementary Table 2).** To identify biological processes most significantly influenced by *Inpp4b*-deficiency, GSEA **(Fig. 4A**) and GO (**Fig. 4B**) were conducted on the differentially expressed gene list (**Supplementary Table 3**). Among several significantly enriched pathways, both methods revealed a highly significant association with the KEGG-lysosome gene set (64 out of 124 genes; corrected *P*-value = 3.62 × 10^-06^). We observed that a disproportionately large number of KEGG lysosomal gene transcripts were downregulated in *Inpp4b^-/-^ MLL-AF9* gene profiles compared to *Inpp4b^+/+^* controls (**Fig. 4C**). Gene expression schematics generated by STRING-db, illustrated gene expression losses were associated with lysosome-active proteins and transporters including several cathepsins (*Ctss, Ctsc and Ctsf*), lysosomal transporters (*Slc11a1*) and other proteases (*Lgmn, Psap, Lipa*) (**Fig. 4C, D**). GSEA using independent lysosomal gene sets curated from the Coordinated Lysosomal Expression and Regulation (CLEAR) network or from the harmonizome **(Fig. 4E-G)** demonstrated similar enrichment profiles from *Inpp4b^-/-^ MLL-AF9* gene profiles compared to *Inpp4b^+/+^* controls (Rouillard et al., 2016; Sardiello et al., 2009). Furthermore, to support our observations that *Inpp4b* deficiency is associated with altered stemness and differentiation phenotypes (**Fig. 3**), we also interrogated genes and networks associated with leukemic stemness or myeloid differentiation. GSEA revealed significant changes in the LSC17 gene signature, a myeloid cell development signature, and an osteoclast differentiation signature in *Inpp4b*-deficiency (**Supplementary Fig 4A-D**). These findings indicate that *Inpp4b*-deficiency in *MLL-AF9* leukemia significantly alters gene transcription networks associated with lysosome biology, myeloid stemness and myeloid lineage differentiation.

**Figure 4.**
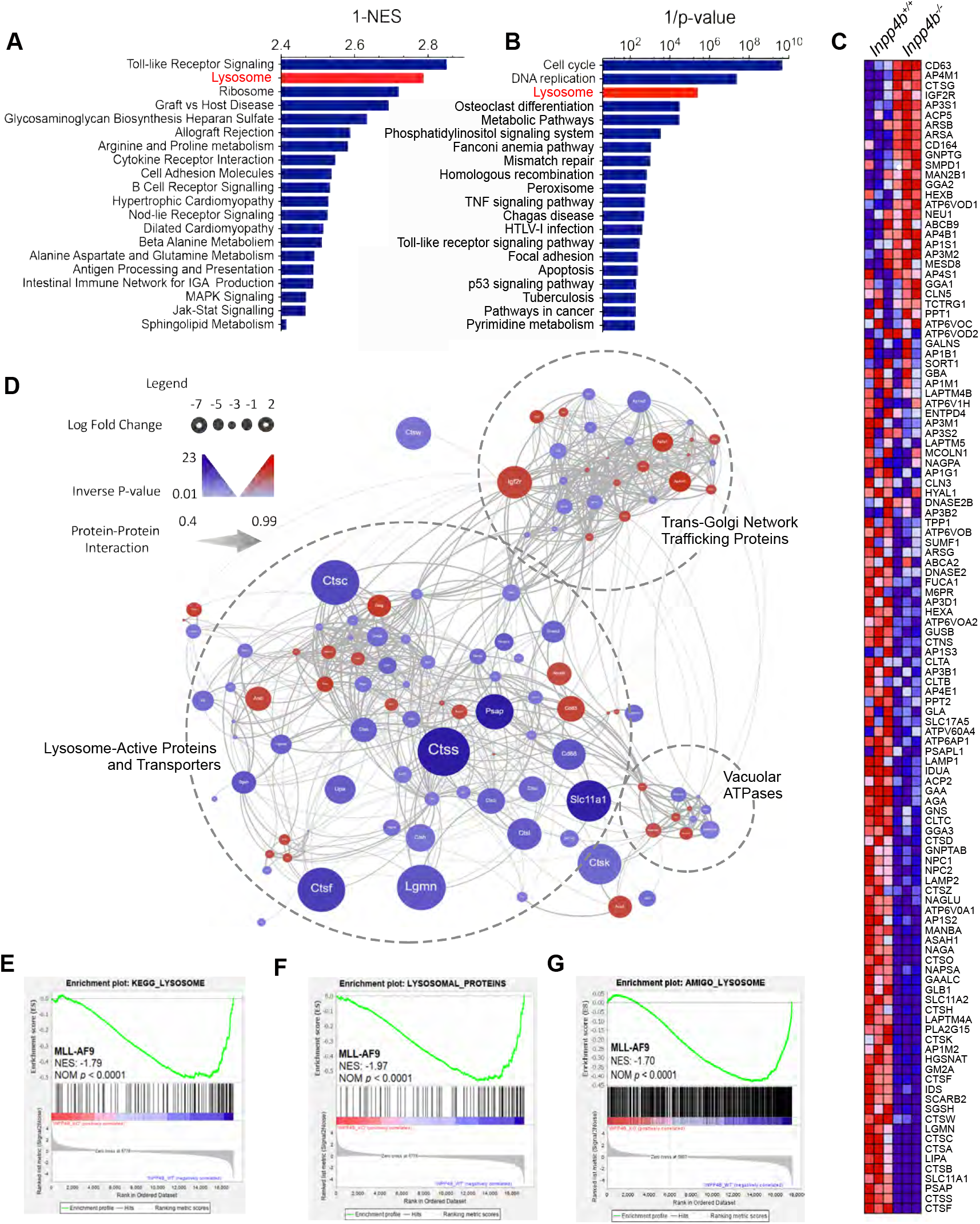
Inpp4b expression is associated with lysosomal gene sets. A. Top 20 enriched KEGG gene sets in Inpp4b+/+ and Inpp4b-/- MLL-AF9 leukemia blast cells determined using GSEA. B. Top 20 enriched KEGG pathways from significantly upregulated genes (log-fold-change > 1, FDR < 0.05) in Inpp4b-/- MLL-AF9 leukemia blast cells were determined using the DAVID analysis tool. C. Heat map of differentially expressed KEGG Lysosome genes from Inpp4b+/+ and Inpp4b-/- MLL-AF9 leukemia blast cells (upregulated in red, downregulated in blue). D. STRING-db analysis of differential expression of KEGG Lysosome genes from Inpp4b+/+ and Inpp4b-/- MLL-AF9 leukemia blast cells. E. Gene set expression analysis (GSEA) for Lysosome pathway genes in Inpp4b-/- MLL-AF9 leukemia blast cells witt the KEGG Lysosome gene set; (F) Lysosomal proteins gene set; (G) Harmonizome (Amigo) lysosome gene set.

### *INPP4B* expression in human AML is associated with lysosomal gene signatures

We identified the top 20 most enriched KEGG-defined gene sets associated with *INPP4B* expression by interrogating AML patient datasets using GSEA (**Fig 5A-D, Supplementary Fig 5A, B)**. Notably, the lysosomal gene set, and other lysosome-related gene sets including glycosaminoglycan degradation and other glycan degradation were ranked among the top 20 most significantly enriched (**Fig 5A-D, Supplementary Fig 5A, B)**. Enrichment scores for the KEGG lysosomal pathways were observed to be significant in TCGA, Valk, Verhaak, and Wouters AML patient datasets (**Fig 5E-H**) (Cancer Genome Atlas Research Network et al., 2013; Herold et al., 2018; Klein et al., 2009; Metzeler et al., 2008; Palmieri et al., 2011; Valk et al., 2004; Verhaak et al., 2009; Wouters et al., 2009). Furthermore, among the 121 genes which comprise the KEGG Lysosome gene set, 78 genes were classified as leading-edge genes (LEGs) in at least one of the TCGA, Verhaak, Valk and Wouters datasets, and a core set of 30 LEGs were present in all 4 datasets (**Fig 5I, J**). In support of our mouse data, analysis of public AML datasets provide further evidence that INPP4B plays a role in regulating lysosomal genes in human AML.

**Figure 5:**
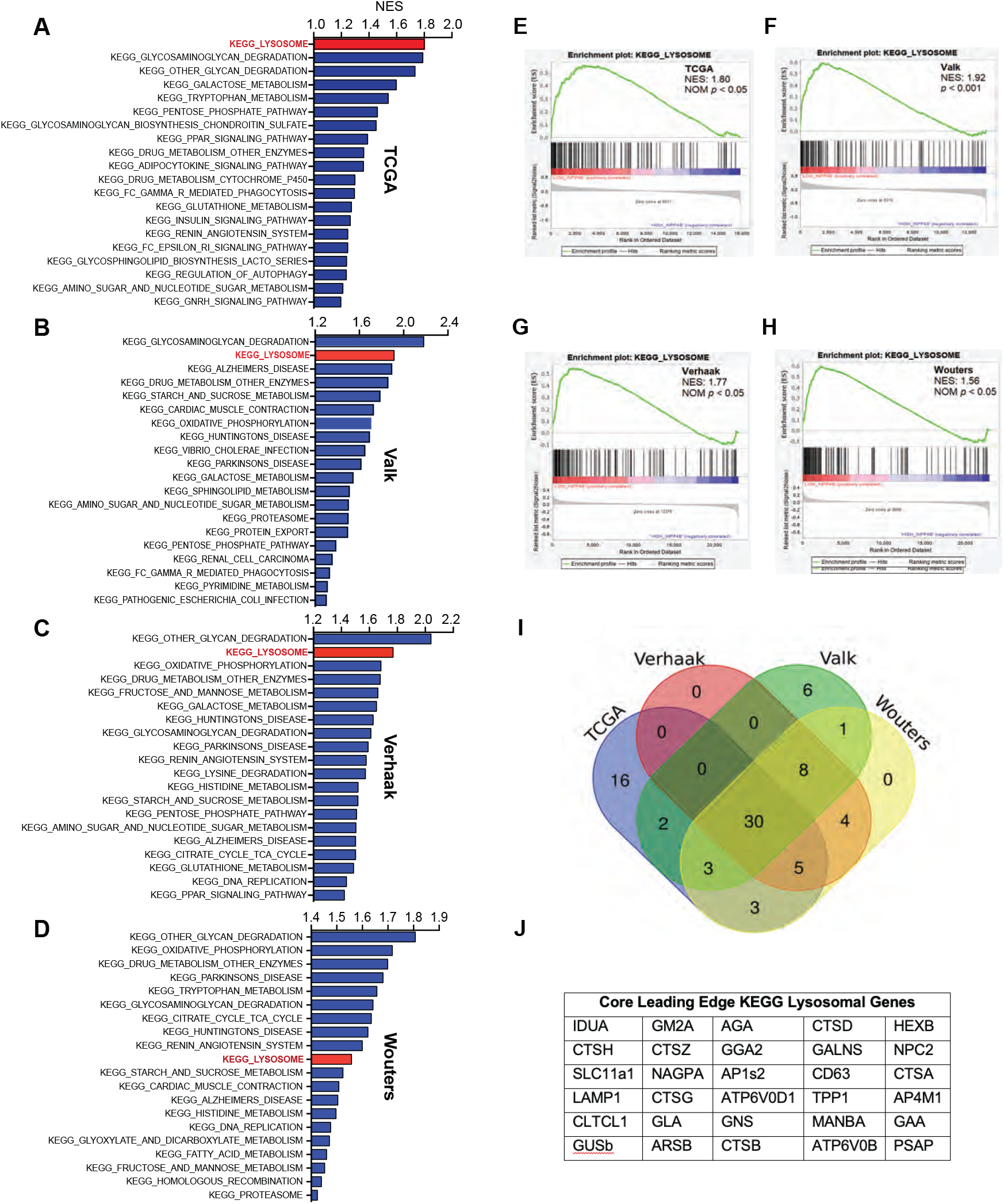
A Lysosome gene set is enriched in patients with low levels of INPP4B. Histograms of the top 20 enriched KEGG gene sets in INPP4B-low patients from the (A) TCGA, (B) Valk (C) Verhaak and (D) Wouters AML patient datasets ranked by normalized enrichment score (NES). GSEA enrichment plots of KEGG Lysosome geneset demonstrating significant enrichment with INPP4B expression in (E) TCGA, (F) Valk (G) Verhaak and (H) Wouters AML patient datasets. I. Venn diagram illustrating the number of genes from the KEGG lysosome gene set that contribute to the leading-edge subset in the four AML datasets. J. Table of the 30 common core enriched lysosome genes among the four AML datasets.

### INPP4B promotes lysosomal gene expression without increasing nuclear translocation of TFEB

To identify a direct and causative role for INPP4B in lysosome biogenesis, we measured the effects of INPP4B overexpression on the levels of several core leading edge KEGG lysosomal genes **(Fig 5J)**. We confirmed that *LAMP1, CTSB, CTSD, ATP6V1H, ATP6V1D*, and *MCOLN1* transcripts were significantly elevated in *INPP4B* overexpressing OCI-AML2 cells **(Fig 6A)**. The promoter region of these and many additional lysosomal genes share the CLEAR consensus promoter element that is targeted by transcription factor EB (TFEB) (Sardiello et al., 2009). We confirmed that *INPP4B* can positively induce lysosomal gene expression using a CLEAR reporter construct, where we observed a significant elevation of luciferase levels **(Fig 6B)**. Thus we hypothesized that *INPP4B* may promote TFEB activity upstream of CLEAR activation. Surprisingly, INPP4B overexpression or knockout in OCI-AML2 did not alter the extent of TFEB nuclear localization in steady state, starved or serum restimulated conditions **(Fig 6C-F and Supplementary Fig 6A-C).** These data show indicate that INPP4B promotes lysosomal biogenesis, however it does so without influencing TFEB nuclear translocation. Further studies are required to elucidate the molecular interaction between INPP4B and lysosome gene expression activation.

**Figure 6:**
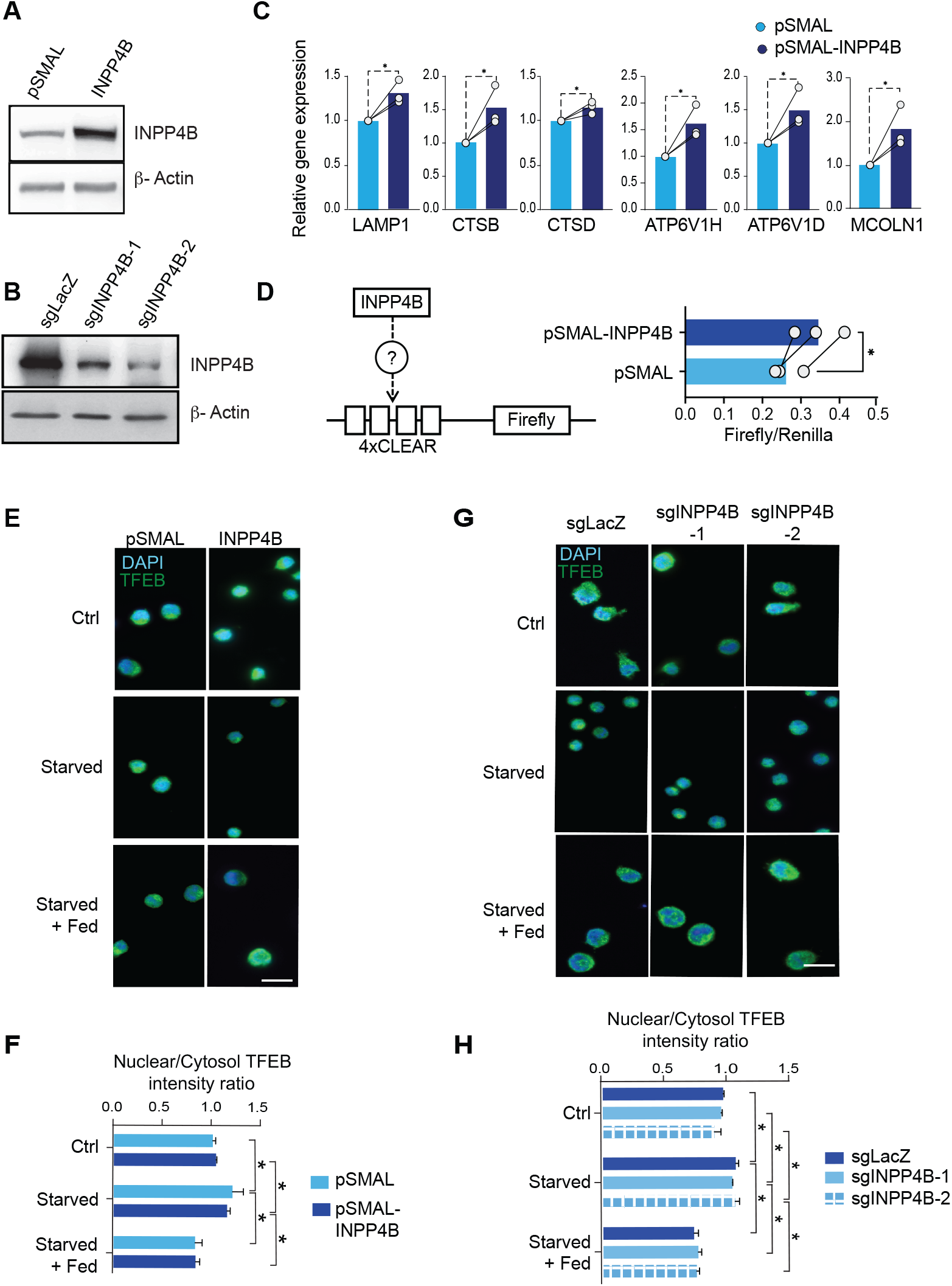
INPP4B promotes expression of lysosomal genes. Representative Western blots demonstrating (A) overexpression and (B) knockdown of INPP4B protein levels in OCI-AML2 cells. C. qRT-PCR analysis of lysosome gene expression in OCI-AML2 cells normalized against Actb. Shown is mean + SEM from three independent experiments. D. HEK293T cells overexpressing control or INPP4B constructs were co-transfected with pGL3 4X-CLEAR-Firefly reporter and CMV-promoter Renilla luciferase internal control and quantitated as the ratio of Firefly to Renilla. E,G. OCI-AML2 cells in the steady state, FCS starved, or starved and FCS fed conditions were immunostained for TFEB and (F,H) analyzed for nuclear/cytosol TFEB intensity.

### INPP4B controls lysosomal content, function and sensitivity to lysosomal inhibition in AML cells

We next investigated a direct role for INPP4B in lysosome biology by evaluating *INPP4B* overexpression or knockout on lysosomal characteristics in AML cells. We measured lysosome levels using Lysotracker Red (LTR) staining in AML cells. *INPP4B*-overexpression produced elevated levels of LTR fluorescence (**Fig. 7A**), whereas knockdown of *INPP4B* expression led to lower levels of LTR fluorescence (**Fig. 7B**). To measure of lysosomal function, we assessed the proteolytic activity of lysosomes in our in OCI-AML2 cell models. Firstly, cathepsin B activity was measured using Magic Red, fluorophore substrate that becomes fluorescent when cleaved by cathepsin B in the lysosome. *INPP4B* overexpressing cells displayed significantly greater Magic Red fluorescence levels, whereas knockout of *INPP4B* reduced Magic Red fluorescence (**Fig. 7C,D**). Additionally, we used DQ-BSA-Green, a sensor that is endocytosed and carried as cargo to lysosomes, where its bright fluorescence is activated by the acidic and highly proteolytic environment (Marwaha and Sharma, 2017). *INPP4B* overexpressing OCI-AML2 cells incubated with DQ-BSA-Green activated significantly greater levels of DQ-BSA-Green fluorescence compared to controls at each time point assessed (**Fig. 7E**). Likewise, experiments conducted in *Inpp4b^+/+^* and *Inpp4b^-/-^ MLL-AF9* leukemias for up to 8 hours showed that *Inpp4b^-/-^* leukemia cells had significantly lower levels of DQ-BSA-Green fluorescence throughout the time-course (**Fig. 7F**). Control experiments using fluorescent-tagged Dextran-Red to measure endocytosis or trafficking rates demonstrated no difference in *INPP4B*-overexpressing or control OCI-AML2 cells, nor *Inpp4b^+/+^* and *Inpp4b^-/-^ MLL-AF9* leukemia cells (**Supplementary Fig 7A-B**).

**Figure 7:**
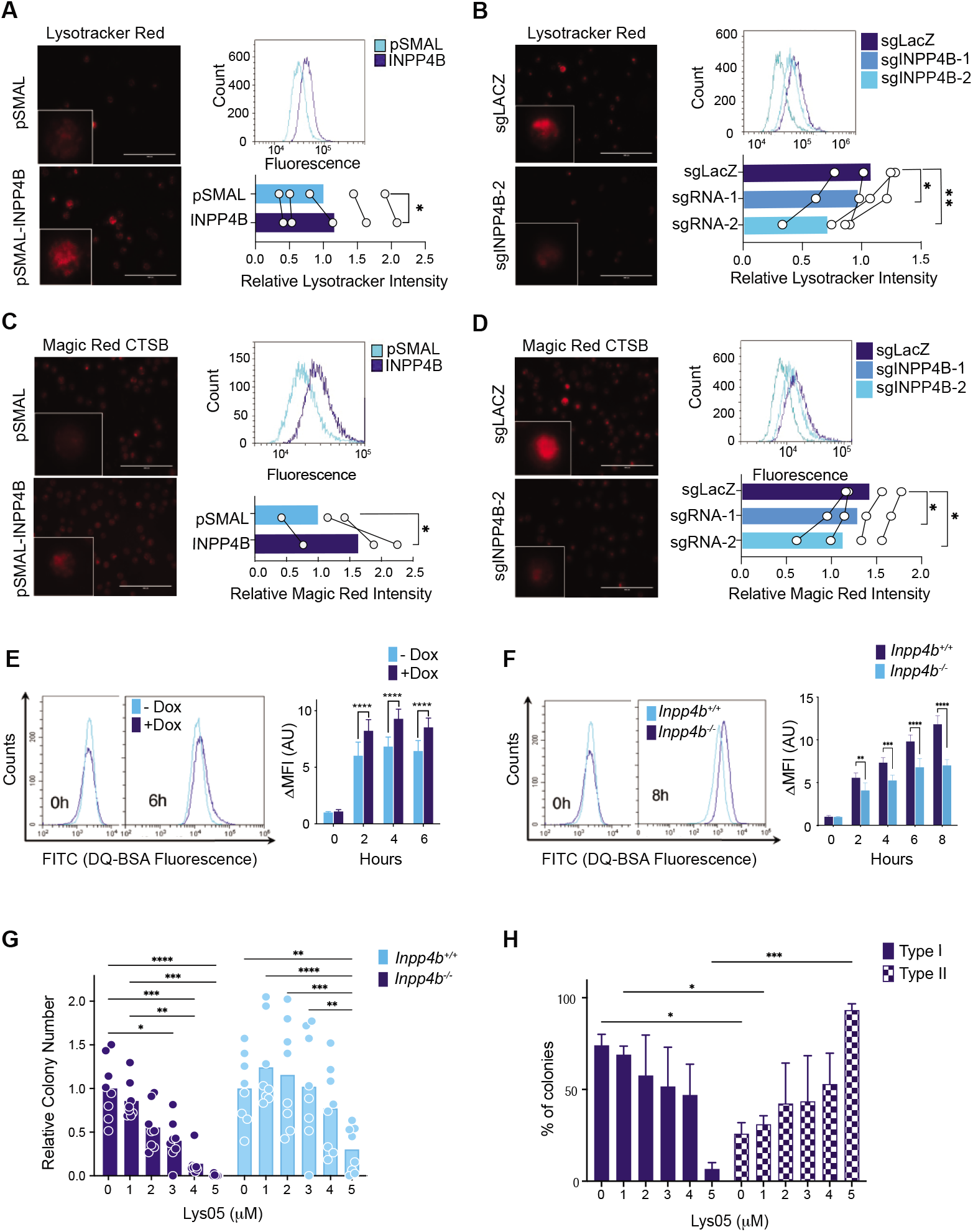
INPP4B regulates lysosomal biology and functions in leukemia cells. INPP4B overexpressing (A) and INPP4B knockout (B) OCI-AML2 cells were labelled with 1uM lysotracker red were imaged by microscopy and quantitated using flow cytometry. INPP4B overexpressing (C) and INPP4B knockout (D) OCI-AML2 cells were labelled with Magic Red-cathepsin B were imaged by microscopy and quantitated using flow cytometry. Scale bar: 15 μm. E. Inducible-INPP4B OCI-AML2 cells were treated without or with doxycycline (100mg/mL) and incubated with DQ-BSA-Green™ and fluorescence was monitored by flow cytometry hourly, for up to 6 hours. Representative flow cytometry histograms displayed. F. Inpp4b+/+ and Inpp4B-/- MLL-AF9 leukemia blasts were incubated with DQ-BSA-Green™ and fluorescence was monitored by flow cytometry hourly, for up to 8 hours. Representative flow cytometry histograms displayed. G. Inpp4b+/+ and Inpp4B-/- MLL-AF9 leukemia blasts were plated in the LFC assay in the presence of Lys05 at indicated dose for 7 days and monitored for total colony counts or (H) percentage of type I and type II colonies. Inpp4b+/+ MLL-AF9 leukemia blasts were plated in the LFC assay in the presence of Lys05 at indicated dose for 7 days. Data represent + SEM from at least three independent triplicate experiments with at least 10,000 events per condition. Students t-test or ANOVA and post-hoc analysis are displayed with *P < 0.05, ** P < 0.01, *** P < 0.001, **** P < 0.0001 compared to indicated control condition.

To further confirm a role for INPP4B in regulating lysosomal properties, we examined the consequences of lysosomal inhibition using the potent lysosomotropic agent Lys05 (Amaravadi and Winkler, 2012; Cechakova et al., 2019; McAfee et al., 2012) in *Inpp4b^+/+^* and *Inpp4b^-/-^ MLL-AF9* leukemia cells. A dose-response assessment in *MLL-AF9* leukemia cells *in vitro* demonstrated that *Inpp4b^+/+^* and *Inpp4b^-/-^* cells had similar sensitivity to the toxic effects of Lys05 (**Supplementary Fig 7C**). To specifically test the effects of Lys05 on leukemic phenotypes, colony forming capacity and colony differentiation were performed using sub-toxic concentrations of Lys05. LFC assays showed that Lys05 reduced the colony forming potential of *Inpp4b^+/+^ MLL-AF9* leukemia cells in a dose-dependent manner. This effect was significantly blunted in *Inpp4b-*deficient cells indicating that lysosomal inhibition reduces the colony forming capacity of *MLL-AF9* leukemia cells in an Inpp4b-dependent manner (**Fig. 7G)**. We reasoned that this may be due to a differentiating effect of lysosomal inhibition on AML blasts. Thus, we performed independent LFC assays in Lys05-treated *Inpp4b^+/+^ MLL-AF9* leukemia cells and specifically recorded the percentage of type I and type II colonies as a measure of colony differentiation status. We observed that Lys05 treatment reduced the percentage of the primitive type-I colonies and concomitantly increased the percentage of more differentiated type-II colonies in a dose dependent manner (**Fig. 7H)**. Together, these results indicate that INPP4B expression increases lysosomal content, promotes lysosomal function, and is necessary for the observed reduction in colony forming capacity and increased differentiation associated with lysosomotropic agents. Together our results suggest a model in which INPP4B promotes the development of AML with poor clinical characteristics by driving lysosomal content and function, which are vital to maintain leukemic stemness and prevent leukemic differentiation (**Supplementary Fig 8**).

## Discussion

By interrogating human AML patient databases, investigating consequences of *Inpp4b*-deficiency in murine leukemogenesis, combined with validation in AML cell line models, we have discovered that *INPP4B* is associated with leukemic stemness; is enriched in LSC; promotes leukemia initiating potential. Furthermore INPP4B regulates lysosome gene transcript expression and lysosomal functions in AML. Finally disruption of lysosome functions by either INPP4B depletion or direct pharmacological inhibition leads to leukemic differentiation (**Supplementary Fig 8**). Overall, by delineating INPP4B functions in AML, we have unveiled a novel INPP4B-lysosome signalling axis that drives ‘stemness’ in leukemia.

By profiling *INPP4B* expression in purified leukemia cell populations and AML datasets, we revealed that *INPP4B* expression was associated with LSC stemness regulation and differentiation. This notion was tested and confirmed in *Inpp4b^-/-^* leukemias which demonstrated a differentiated AML phenotype and reduced leukemia initiating potential. The association with leukemic stemness identified INPP4B as a putative therapeutic target, with potential to compromise LSC self-renewal function and consequently drive differentiation of AML. However, given that INPP4B inhibition is not currently feasible, investigated processes downstream of INPP4B signaling with pharmacological potential. Expression profiling of *Inpp4b*-deficeint leukemias combined with gene network analyses revealed that the expression of key lysosomal genes including proteases and membrane transporters were strongly associated with *INPP4B* status. This finding was corroborated in AML patient data and may represent rational candidate drug targets for AML therapy.

Long known as organelles that serve as a cell’s main degradative centre (Luzio et al., 2000; Settembre and Ballabio, 2014), lysosomes are now recognized as as key cellular sentinels of nutrient concentration and metabolic activity with critical roles in integrating diverse signal transduction pathways (Inpanathan and Botelho, 2019; Lamming and Bar-Peled, 2019; Lawrence and Zoncu, 2019; Savini et al., 2019). Crucially, studies implicating lysosomal functions in stem cell maintenance are emerging. For instance, functional lysosomes in neuronal stem cells are needed to mitigate protein-aggregate accumulation, a consequence that would otherwise promote aberrant differentiation, increase aging, and reduce stemness (Leeman et al., 2018). Lysosomes are crucial for the exquisite regulation of cell cycling required in quiescent hematopoietic stem cells (HSC) and they are asymmetrically distributed during division of stem cells, and thus predictive of future daughter cell fates (Loeffler et al., 2019). Lysosomes and also impact HSC fate through their ability to integrate diverse cellular cues and coordinate diverse signal transduction pathways (García-Prat et al., 2021). In leukemia, key lysosomal signaling networks have been implicated in fate and differentiation (Yun et al., 2021), and increased lysosomal mass and biogenesis in leukemic progenitor cells selectively sensitize AML to lysosomal inhibitors (Bernard et al., 2015; Sukhai et al., 2013). Our work adds to this increasing body of knowledge by identifying upstream regulatory pathways that can modulate lysosomal activity and thereby regulate AML stemness.

Rogers *et al*. recently showed that INPP4B directly enhances late endosome/lysosome formation and drives cell proliferation and tumor growth in *PIK3CA*-mutant ER^+^ breast cancers (Rodgers et al., 2021). This was attributed to a mechanism whereby INPP4B-mediated PI(3,4)P2 to PI(3)P conversion on late endosomes promotes Wnt/β-catenin signaling. Notably, aberrant Wnt/β-catenin signaling has been linked with adverse clinical outcomes in AML (Frenquelli and Tonon, 2020; Grainger et al., 2018). In our studies, leukemic cell models with INPP4B overexpression and INPP4B depletion demonstrated that INPP4B can regulate lysosomal homeostasis through regulation of lysosomal gene expression and thereby increase lysosomal content, and the proteolytic capacity of lysosomes. Together, the study by Rogers *et al*. and our work indicate that INPP4B may have both direct and indirect roles in promoting lysosome content and function, respectively.

Lysosomes have emerged as an intriguing organelle with effects on the initiation, progression and therapy of cancer. It is clear that lysosomal functions can contribute to several cancer hallmarks, including proliferation, altered metabolism, invasion and metastasis among others (Rafiq et al., 2021). In leukemia and hematopoiesis, lysosomes can influence stemness and differentiation, a newly proposed cancer hallmark (Hanahan, 2022). Critically, lysosomes have also been implicated in chemoresistance through a lysosomal sequestration mechanism which is triggered by transcriptional lysosomal biogenesis (Zhitomirsky and Assaraf, 2016). Given their pleiotropic roles in cancer cell biology, lysosomes have become an attractive target for cancer therapy (Rafiq et al., 2021). Our observations with INPP4B in AML are resonant with the activation of transcriptional lysosome biogenesis which promotes cancer progression, restricted leukemic differentiation and chemoresistance. Thus it is tempting to propose that lysosomal inhibition may compromise AML by promoting LSC differentiation and simultaneously combat chemotherapy resistance. Moreover, in addition its ability to predict poor disease in AML, *INPP4B* expression may also serve as a biomarker of response to lysosomal inhibition in line with the notion that lysosomes in AML cells are bigger in size which could make AML cells more susceptible towards lysosomal disruption than normal cells (Bernard et al., 2015; Sukhai et al., 2013)

## STAR Methods

### Key reagents Table

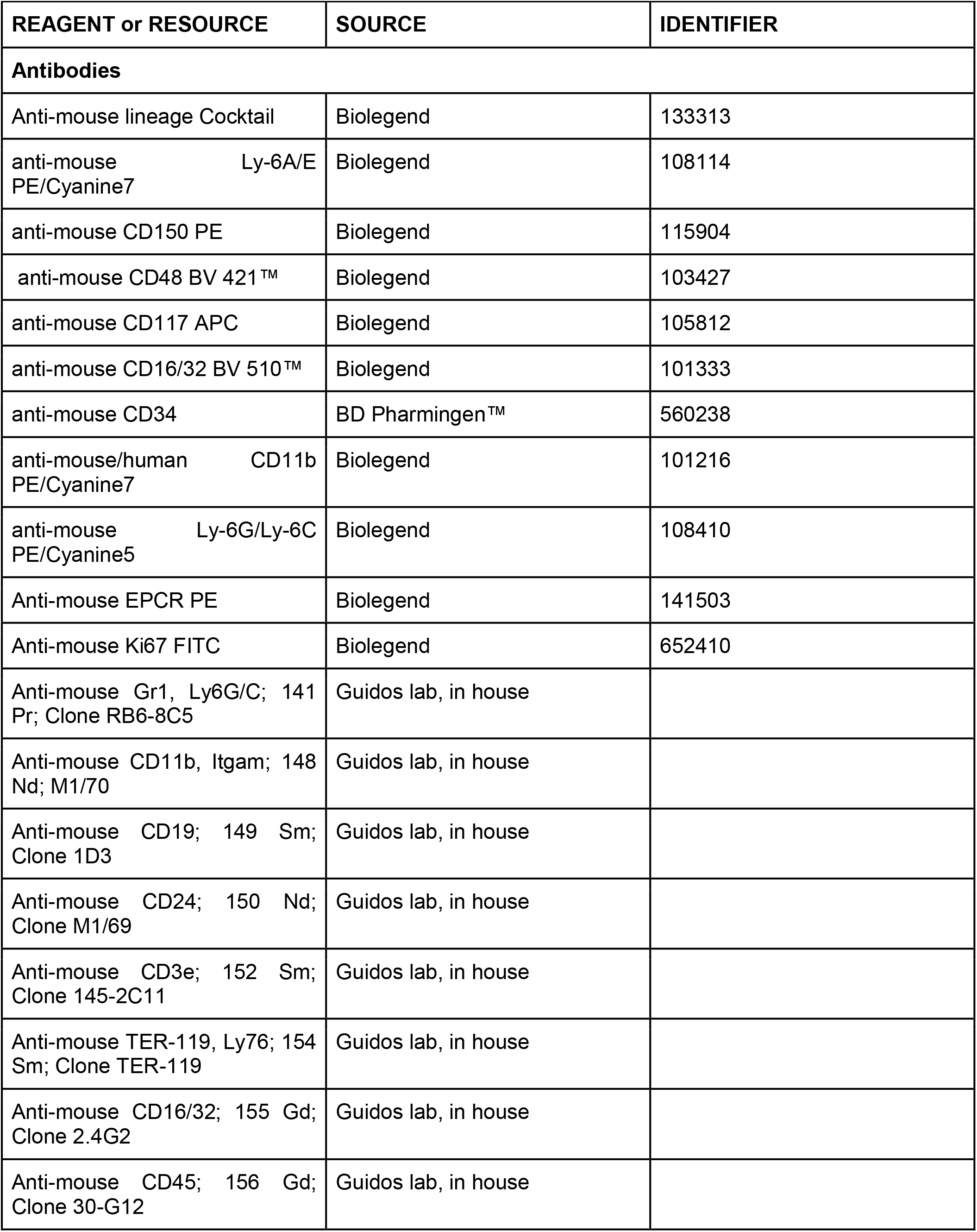
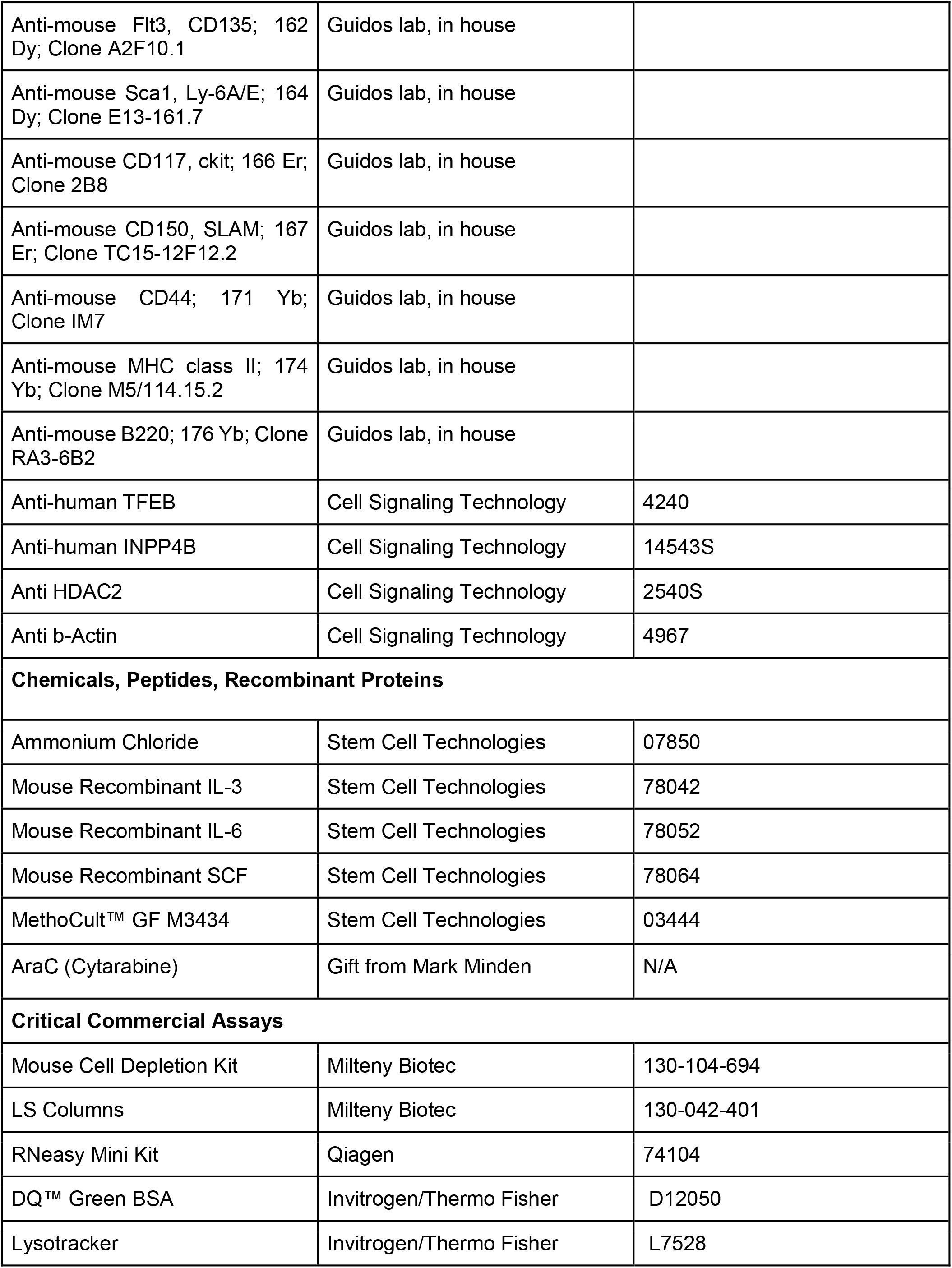
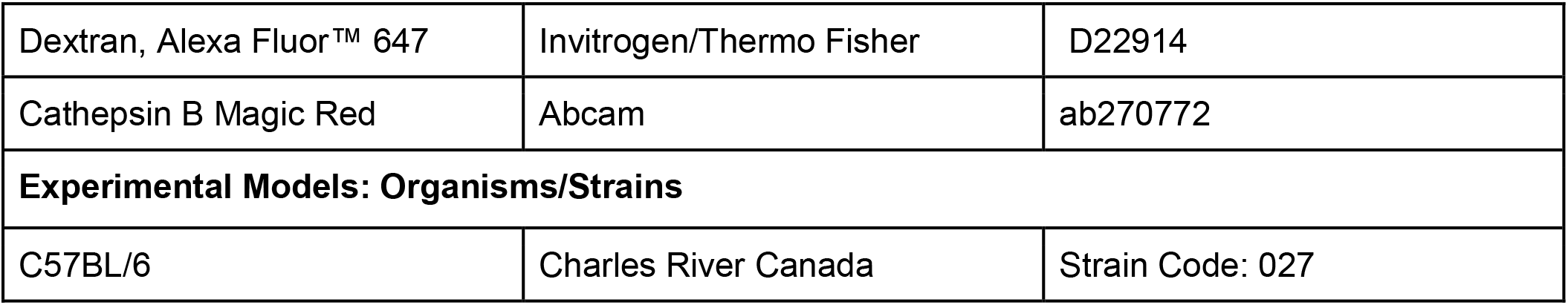

| REAGENT or RESOURCE | SOURCE | IDENTIFIER |
| --- | --- | --- |
| **Antibodies** |  |  |
| Anti-mouse lineage Cocktail | Biolegend | 133313 |
| anti-mouse Ly-6A/E<br>PE/Cyanine7 | Biolegend | 108114 |
| anti-mouse CD150 PE | Biolegend | 115904 |
| anti-mouse CD48 BV 421™ | Biolegend | 103427 |
| anti-mouse CD117 APC | Biolegend | 105812 |
| anti-mouse CD16/32 BV 510™ | Biolegend | 101333 |
| anti-mouse CD34 | BD Pharmingen™ | 560238 |
| anti-mouse/human CD11b<br>PE/Cyanine7 | Biolegend | 101216 |
| anti-mouse Ly-6G/Ly-6C<br>PE/Cyanine5 | Biolegend | 108410 |
| Anti-mouse EPCR PE | Biolegend | 141503 |
| Anti-mouse Ki67 FITC | Biolegend | 652410 |
| Anti-mouse Gr1, Ly6G/C; 141<br>Pr; Clone RB6-8C5 | Guidos lab, in house |  |
| Anti-mouse CD11b, Itgam; 148<br>Nd; M1/70 | Guidos lab, in house |  |
| Anti-mouse CD19; 149 Sm;<br>Clone 1D3 | Guidos lab, in house |  |
| Anti-mouse CD24; 150 Nd;<br>Clone M1/69 | Guidos lab, in house |  |
| Anti-mouse CD3e; 152 Sm;<br>Clone 145-2C11 | Guidos lab, in house |  |
| Anti-mouse TER-119, Ly76; 154<br>Sm; Clone TER-119 | Guidos lab, in house |  |
| Anti-mouse CD16/32; 155 Gd;<br>Clone 2.4G2 | Guidos lab, in house |  |
| Anti-mouse CD45; 156 Gd;<br>Clone 30-G12 | Guidos lab, in house |  |
| Anti-mouse Flt3, CD135; 162 Dy; Clone A2F10.1 | Guidos lab, in house |  |
| Anti-mouse Sca1, Ly-6A/E; 164 Dy; Clone E13-161.7 | Guidos lab, in house |  |
| Anti-mouse CD117, ckit; 166 Er; Clone 2B8 | Guidos lab, in house |  |
| Anti-mouse CD150, SLAMF; 167 Er; Clone TC15-12F12.2 | Guidos lab, in house |  |
| Anti-mouse CD44; 171 Yb; Clone IM7 | Guidos lab, in house |  |
| Anti-mouse MHC class II; 174 Yb; Clone M5/114.15.2 | Guidos lab, in house |  |
| Anti-mouse B220; 176 Yb; Clone RA3-6B2 | Guidos lab, in house |  |
| Anti-human TFEB | Cell Signaling Technology | 4240 |
| Anti-human INPP4B | Cell Signaling Technology | 14543S |
| Anti HDAC2 | Cell Signaling Technology | 2540S |
| Anti b-Actin | Cell Signaling Technology | 4967 |
| **Chemicals, Peptides, Recombinant Proteins** |  |  |
| Ammonium Chloride | Stem Cell Technologies | 07850 |
| Mouse Recombinant IL-3 | Stem Cell Technologies | 78042 |
| Mouse Recombinant IL-6 | Stem Cell Technologies | 78052 |
| Mouse Recombinant SCF | Stem Cell Technologies | 78064 |
| MethoCult™ GF M3434 | Stem Cell Technologies | 03444 |
| AraC (Cytarabine) | Gift from Mark Minden | N/A |
| **Critical Commercial Assays** |  |  |
| Mouse Cell Depletion Kit | Milteny Biotec | 130-104-694 |
| LS Columns | Milteny Biotec | 130-042-401 |
| RNeasy Mini Kit | Qiagen | 74104 |
| DQ™ Green BSA | Invitrogen/Thermo Fisher | D12050 |
| Lysotracker | Invitrogen/Thermo Fisher | L7528 |
| Dextran, Alexa Fluor™ 647 | Invitrogen/Thermo Fisher | D22914 |
| Cathepsin B Magic Red | Abcam | ab270772 |
| **Experimental Models: Organisms/Strains** |  |  |
| C57BL/6 | Charles River Canada | Strain Code: 027 |

#### Gene expression data

Genome-wide expression data from The Cancer Genome Atlas (TCGA)-LAML dataset was downloaded from the ICGC database (https://icgc.org/). Normalized microarray data for the Verhaak (GSE6891), Valk (GSE1159), Herold (37642), Klein (GSE15434), Metzeler I (GSE12417), Metzeler II (GSE12417), Wouters (GSE14468), and Balgobind (GSE17855) databases were downloaded from GEO database (http://www.ncbi.nlm.nih.gov/geo/). GO analysis was performed using The Database for Annotation, Visualization and Integrated Discovery [DAVID; (Huang et al., 2009a)]. The top 200 enriched genes in *INPP4B^high^* patients from the TCGA, Verhaak, Wouters, Herold, Klein and Metzeler datasets were submitted to the DAVID analysis tool to identify enriched GO Biological Processes terms (Huang et al., 2009b, 2009a). Enriched terms related to Phosphatidylinositol signaling or the hematopoietic compartment, and the corresponding datasets were highlighted in a chord diagram that was generated with the circlize R package (Gu et al., 2014).

#### Gene Set Enrichment Analysis (GSEA)

GSEA was performed on the datasets using GSEA v.4.0.3 provided by the Broad Institute (http://software.broadinstitute.org/gsea/downloads.jsp). Samples were rank ordered and split by *INPP4B* status, 25% high/75% low (human) or *Inpp4b^+/+^/Inpp4b^-/-^* (mouse). Enriched gene sets were identified by 1,000 phenotype permutations in the human datasets, and 1,000 gene set permutations in the mouse dataset. Gene sets with a nominal *p*--value < 0.05 were considered significantly enriched. The curated KEGG (CP: KEGG) and BROWN_MYLEOID_CELL_DEVELOPMENT_UP gene set were obtained from MSigDB Collections (http://software.broadinstitute.org/gsea/msigdb/collections.jsp) and the LSC17 and Lysosomal Proteins gene sets were generated from the indicated publication (Ng et al., 2016; Palmieri et al., 2011).

#### Gene Network Analysis

The lysosome interacting genes were passed through String-db.v.11 (Szklarczyk et al., 2019). and the combined binding score for each protein-protein interaction was obtained. This output, which is experimentally determined and annotated, connects the protein nodes. The absolute log fold change from the RNAseq is used to size the nodes and the colour corresponds to the -log10(Pvalue). The interaction map was generated by ‘ForceAtlas2’ in Gephi.v.0.9.2. (Jacomy et al., 2014).

#### Mice

*Inpp4b^-/-^* mice kindly provided by Jean Vacher (Institut de Recherches Cliniques de Montréal) and were crossed into the C57BL/6 and genotyped as previously described (Ferron et al., 2011). C57BL/6 wild-type mice were bred and housed at the University Health Network/Princess Margaret Cancer Center and the Division of Comparative Medicine Facility at the University of Toronto. B6.SJL-Ptprca Pepcb/BoyJ (CD45.1) mice were purchased from The Jackson Laboratory (Bar Harbor, Maine, USA). All animal experiments were performed in accordance with national and institutional guidelines approved by the Canadian Counsel on Animal Care and approved by the University Health Network Animal Care Committee and the Division of Comparative Medicine at the University of Toronto.

#### *MLL-AF9* AML Models

*MLL-AF9* expressing cells were generated by retroviral transduction of murine bone marrow progenitors. Briefly, bone marrow was flushed from the long bones of 10-week-old *Inpp4b^+/+^* and *Inpp4b^-/-^* mice. From these total bone marrow preparations, 2 x 10^6^ LSK cells were sorted and grown for 24 hours in IMDM supplemented with 20 ng/ml SCF, 10 ng/ml IL-6 and 10 ng/ml IL-3 (Gibco). *Inpp4b^+/+^* and *Inpp4b^-/-^* LSK were each retrovirally transduced with pMSCV-*MLL-AF9*-IRES-mVenus. 24 hours post retroviral transduction, 2 x 10^5^ Venus^+^ cells were transplanted to sub-lethally irradiated (4.5Gy) donor C57BL/6 mice to generate primary leukemias (Zuber et al., 2009). Subsequent transplantation of mVenus^+^ cells from mice with terminal leukemias were performed with 250 cells and designated as secondary leukemia assays. *In vitro* leukemia forming cell (LFC) assays were performed in IMDM medium with 2% methylcellulose, supplemented with 20 ng/ml SCF, 10 ng/ml IL-6 and 10 ng/ml IL-3 (Stem Cell Technologies). Cytological analysis of blood and bone marrow smears from *Inpp4b^+/+^* and *Inpp4b^-/-^* leukemic mice was done by Wright-Giemsa staining (Sigma). For limiting dilution transplantation assays (LDA), *MLL-AF9* leukemia cells were injected at defined doses (equivalent to 10, 25, 50, 100, and 1000 blast cells) into 8-week-old female C57BL/6 mice. LSC frequency was estimated using the online tool ELDA (http://bioinf.wehi.edu.au/software/elda/index.html) (Hu and Smyth, 2009). AraC (or saline vehicle control) was delivered i.p. at 100 mg/kg for 7 days starting at day 14 post-transplant in mice transplanted with 1000 *MLL-AF9* leukemia cells of indicated genotype.

#### RNA sequencing

*Inpp4b^+/+^* or *Inpp4b^-/-^ MLL-AF9* leukemias (3 in total for each genotype) were sorted for Venus+ before isolation of total cellular RNA with a RNeasy isolation kit (Qiagen). A Bioanalyzer 2100 (Agilent) was used for quality control and quantification. Illumina MouseRef-8 v2.0 Expression BeadChip kits were used for genome-wide expression profiling according to standard protocols at The Centre for Applied Genomics core facility at the Hospital for Sick Children. R Bioconductor 2.13.0 software was used for data processing and other statistical analyses. Raw signals from 25,697 probes were preprocessed for background subtraction, quantile normalization and log2 transformation before the use of moderated t-tests from the Bioconductor software package Limma (linear models for microarray data). Empirical Bayes smoothing was applied to the standard errors. Paired t-tests were used for the identification of differentially expressed genes expression in each genotype subset, and the false-discovery rate (FDR) was estimated with the Benjamini-Hochberg method to correct for multiple testing. Pearson correlations showed that technical replicates had very high correlations between chips. For genes represented by multiple probe sets on the array, we selected the ones with the highest ANOVA F-statistics (lowest FDR-adjusted q value).

#### Mass Cytometry

Purified mAbs were conjugated with heavy metals by the Flow and Mass Cytometry Facility, The Hospital for Sick Children. Murine AML blasts were counted and 2 x10^6^ cells for each sample were stained for cell surface markers in staining media (PBS containing 1% BSA and 0.02% NaN3) for 30 minutes at 4C. Cells were washed with protein-free PBS, stained with 1 mmol/L cisplatin for 5 minutes at room temperature, fixed using the transcription factor buffer set (BD Biosciences) followed by intracellular staining for 60 minutes at 4C. Cells were washed with staining media and stained with 100 nmol/L iridium-labeled DNA-intercalator (Fluidigm) in PBS containing 0.3% saponin and 1.6% formaldehyde at 4 C for up to 48 hours. Cells were washed twice with deionized water prior to adding EQ normalization beads containing Ce140, Eu151, Eu153, Ho165, and Lu175 (Fluidigm) and acquiring on a Helios mass cytometer by The Flow and Mass Cytometry Facility, The Hospital for Sick Children. After normalizing and randomizing values near zero using the Helios software, FCS files were uploaded to Cytobank for analysis.

#### Cells culture and cellular assays

*MLL-AF9* leukemia cells were maintained in IMDM medium supplemented with 10% fetal bovine serum and cytokines: 20 ng/ml SCF, 10 ng/ml IL-6 and 10 ng/ml IL-3 (Stem Cell Technologies). OCI-AML2 cells were cultured in alpha minimal essential medium with 10% fetal bovine serum, 100 units/ml penicillin and 100 ug/ml streptomycin at 37 °C and 5% CO2. Lentiviruses were produced by calcium phosphate transfection of 293T cells with psPAX2 and VSVG as described by the manufacturer (Life Technologies, Burlington, ON, Canada). Viral particles in supernatants collected at 48 and 72 h post-transfection were enriched with Lenti-X Concentrator (Clontech, Mountain View, CA, USA). OCI-AML2 were infected with lentivirus for 24 h with 8 ug/ml protamine sulfate and selected with 2ug/mL puromycin selection. Generation of pSMAL-Puro-FLAG-INPP4B was previously described (Dzneladze et al., 2015). Inducible INPP4B-OCI-AML2 were generated by inserting a donor vector containing an inducible TRE3G-3xFLAG-INPP4B and constitutive Tet-On 3G expression cassettes into the *AAVS1* safe harbor locus by CRISPR editing using an AAVS1-specific guide construct pLCKO_AAVS1_sgRNA_2, a gift from Jason Moffat (Addgene plasmid # 155085; http://n2t.net/addgene:155085; RRID:Addgene_155085). 3xFLAG-INPP4B was cloned by PCR into AscI and PacI sites of the pAAVS1-tet-iCas9-BFP2 vector, a gift from Jason Moffat (Addgene plasmid # 125519; http://n2t.net/addgene:125519; RRID:Addgene_125519). INPP4B was knocked out in constitutive Cas9 expressing OCI-AML2 cells, generated by infecting with lentiCas9-Blast [a gift from Feng Zhang (Addgene plasmid # 52962; http://n2t.net/addgene:52962; RRID:Addgene_52962)] followed by blasticidin selection. Cas9 activity was confirmed using pXPR_011 [(a gift from John Doench & David Root (Addgene plasmid # 59702; http://n2t.net/addgene:59702; RRID:Addgene_59702)]. Lentiviral INPP4B sgRNAs were generated by cloning the following sequences TTTCGCAGATTGTCCCAATG (*INPP4B*-sgRNA-1) and ACTTGTGAACTGACAGTCCC (*INPP4B*-sgRNA-2) into lentiGuide-Puro [a gift from Feng Zhang (Addgene plasmid # 52963; http://n2t.net/addgene:52963; RRID:Addgene_52963)].

#### Luciferase Assay

HEK293T cells were cotransfected with 4X-CLEAR promoter-reporter construct [a gift from Albert La Spada (Addgene plasmid # 66800; http://n2t.net/addgene:66800; RRID:Addgene_66800)] and a CMV-promoter Renilla luciferase vector was used as the internal control with either pSMAL empty vector or pSMAL-INPP4B. Three biological triplicates were performed and displayed as fold increases, statistics were performed by using a students t-test where *P* < 0.05 was significant.

#### Lysosome Assays

For Lysotracker Assays the LysoTracker™ Red DND-99 (Thermo L7528) reagent was used. 0.5 x 10^6^ OCI-AML2 cells were stained in the dark with 1mM Lysotracker in 500 mL of PBS for 15 min at 37°C. For Cathepsin B Assays, the Magic Red Cathepsin B Assay Kit (Abcam ab270772) was used as according to supplied protocol. Briefly, after preparing Magic Red reagent 1 x 10^6^ OCI-AML2 cells in 500 mL were stained in the dark for 30 min at 37°C. After staining of cells with either reagent, cells were washed with PBS and imaging and quantitation were as follows. 1 x 10^5^ stained cells were centrifuged at 200 xg for 10 min onto poly-L-Lysine coated coverslips were immediately mounted onto microscope slides with DAKO fluorescent and imaged with EVOS-FL fluorescent inverted microscope controlled by EVOS XL core imaging system at 20x 0.4 N.A. objective (Thermo Fisher). Remaining cells were analysed by flow cytometry with Beckman Coulter Cytoflex flow cytometer (Beckman, Brea, CO). Background signal was determined with non-labelled cells. A minimum of 10,000 gated events were counted per condition using the phycoerythrin (PE-A) channel for quantitation. For DQ-BSA experiments, OCI-AML2-Tet-ON-INPP4B cells (treated with or without doxycycline (100ng/mL) 48 h prior to lysosome labelling) or MLL-AF9 Inpp4b^+/+^ and Inpp4b^-/-^ cells were incubated with 10ug/mL DQ Red BSA (Invitrogen) or 2uM Alexa-647-conjugated 10kDa dextran (Invitrogen) in complete media for 1h to 6h at 37°C. At each time point, cells were washed twice with PBS and subsequently analyzed for by flow cytometry with Beckman Coulter Cytoflex flow cytometer (Beckman, Brea, CO). A total of 60,000 events were collected per sample per time point using the APC channel. Non-labeled cells were used as control for determining background signal and as time point 0. At least three biological triplicates were performed and displayed as relative fluorescence intensity, statistics were performed by using a paired students t-test where *P* < 0.05 was significant.

#### Immunoblotting and Immunofluorescence

Western blotting was performed with anti-INPP4B (#8450), TFEB (#4240), HDAC2 (#2540) and b-Actin (#4967) from Cell Signaling Technology. For TFEB immunostaining OCI-AML-2 cells were centrifuged at 200xg for 10 min onto poly-L-Lysine coated slides and fixed with 4% paraformaldehyde and permeabilized with 0.1 % Triton X-100 in PBS for 10 min followed by blocking with blocking buffer (10% FBS and 5% BSA in PBS). Cells were incubated overnight at 4°C with mouse anti-human TFEB monoclonal antibody (1:50, R&D systems) in blocking buffer. Dylight-488 conjugated goat anti-mouse IgG secondary antibody (1:1000, Thermo) was added to cells in 0.025% Tween-20 in PBS for 1 h at room temperature. Nuclei were stained with 1 μg/ml DAPI and coverslips were mounted onto microscope slides with DAKO fluorescent mounting media. Cells imaged with ZEISS AxioImager M2 Epifluorescence microscope connected to AxioCam MRm CCD camera and controlled by AxioVision Software Version 4.8 at 40x 1.4 N.A. objective (Carl Zeiss, Oberkochen, Germany). Images were imported onto ImageJ. Intensity threshold applied to select DAPI positive regions and generating a mask, which was applied to green (TFEB immunostain) channel to measure TFEB intensity on DAPI positive regions. Whole cell TFEB intensity was recorded to obtain nuclear to cytosol TFEB intensity ratio.

#### RT-PCR

Qiagen RNeasy mini kit (Qiagen, Venlo, Netherlands) used to extract RNA from OCI-AML2 cells. Superscript IV Vilo cDNA synthesis kit (Thermo) was used to reverse transcribe equal amounts of RNA. TaqMan Fast Advanced Master mix (Applied Biosystems, Foster City, CA) and Taqman assays used to amplify cDNA through quantitative PCR according to manufacturer instructions with QuantStudio 3 Real-Time PCR system (Thermo Fisher) controlled by QuantStudio Design and AnalysisSoftware version 1.2 (Thermo Fisher). Taqman assays include Actb (Hs03023943_g1), CtsD (Hs00924259_m1), CtsB (Hs00157194_m1), Atp6v1h (Hs00977530_m1), Atp6v1d (Hs00211133_m1), Lamp1 (Hs05049889_s1) and Mcoln1 (Hs01100661_g1) and performed in triplicates. Gene expression determined through relative quantification (ΔΔCt method) and normalized to Actb. Three biological triplicates were performed and displayed as fold increases, statistics were performed by using a students t-test where *P* < 0.05 was significant.

#### Statistical Analysis

Errors bars show standard error of the mean (SEM). Data were analyzed with a two-tailed Student t-test for comparison of the means of two groups and by one-way ANOVA unless otherwise stated. *P* values of less than 0.05 were considered statistically significant. No randomization of mice or ‘blinding’ of researchers to sample identity was used during the analyses. Sample sizes were not predetermined on the basis of expected effect size, but rough estimations were made on the basis of pilot experiments and measurements. No data exclusion was applied.

## Acknowledgments

We thank all current and past members of the Salmena and Minden labs for technical assistance and critical discussions. We specifically thank Rod Bremner, Robert C. Laister for critical advice and editing, Norman Iscove for technical counsel; Johannes Zuber for *pMSCV-MLL-AF9-IRES-mVenus* plasmid; Lev Kats for invaluable advice and reagents;. L.S. is the recipient of a Tier II Canada Research Chair (CRC) and was supported through the Human Frontier Career Development Program (HFSP) Award. This work was supported in part by funds from the Department of Pharmacology and Toxicology and Temerty Faculty of Medicine, University of Toronto and awards from Canada Foundation for Innovation (CFI-#33505); The Natural Sciences and Engineering Research Council of Canada (NSERC-RGPIN-2015-03984); Leukemia and Lymphoma Society of Canada (LLSC-Operating Grant #317359; Operating Grant #422332); Acute Leukemia Translational Research Initiative through funding provided by the Ontario Institute for Cancer Research and Government of Ontario; Leukemia Research Foundation (LRF-New Investigator Award #169456), and Canadian Institutes for Health Research (CIHR-Operating Grant #149032; Operating Grant #399716) awarded to L.S. This work was supported in part by funds from CIHR (MOP# 123343) awarded to J.V. Computational analyses were supported in part by funds from NSERC (# 203475), CFI (#225404, #30865), Ontario Research Fund (RDI #34876), IBM and Ian Lawson van Toch Fund awarded to I.J. R.J.B. is a recipient of a Tier II CRC, and this work was supported in part by the NSERC (RGPIN-2015-05489 and RGPIN-2020-043343).

## Authorship Contributions

J.F.W., K.C., G.G., D.K.C.L., G.T.S., C.M.P.M., C.A.N., A.H, CE.M.S., S.Z.X. M.H.S., M.S.G, R.H., A.R. and L.S. performed experiments. D.K.C.L., M.G., M.K., I.D. and I.J. performed bioinformatic analyses. J.V. generated *Inpp4b* mouse models. C.J.G, J.E.D., R.B., and M.D.M. provided critical reagents, mentorship, and expertise. J.F.W. and L.S. conceived the project, planned the study, and wrote the paper. L.S. secured funding for this study. All authors read and approved the text.

## Supplementary Information

**Supplementary Figure 1.**
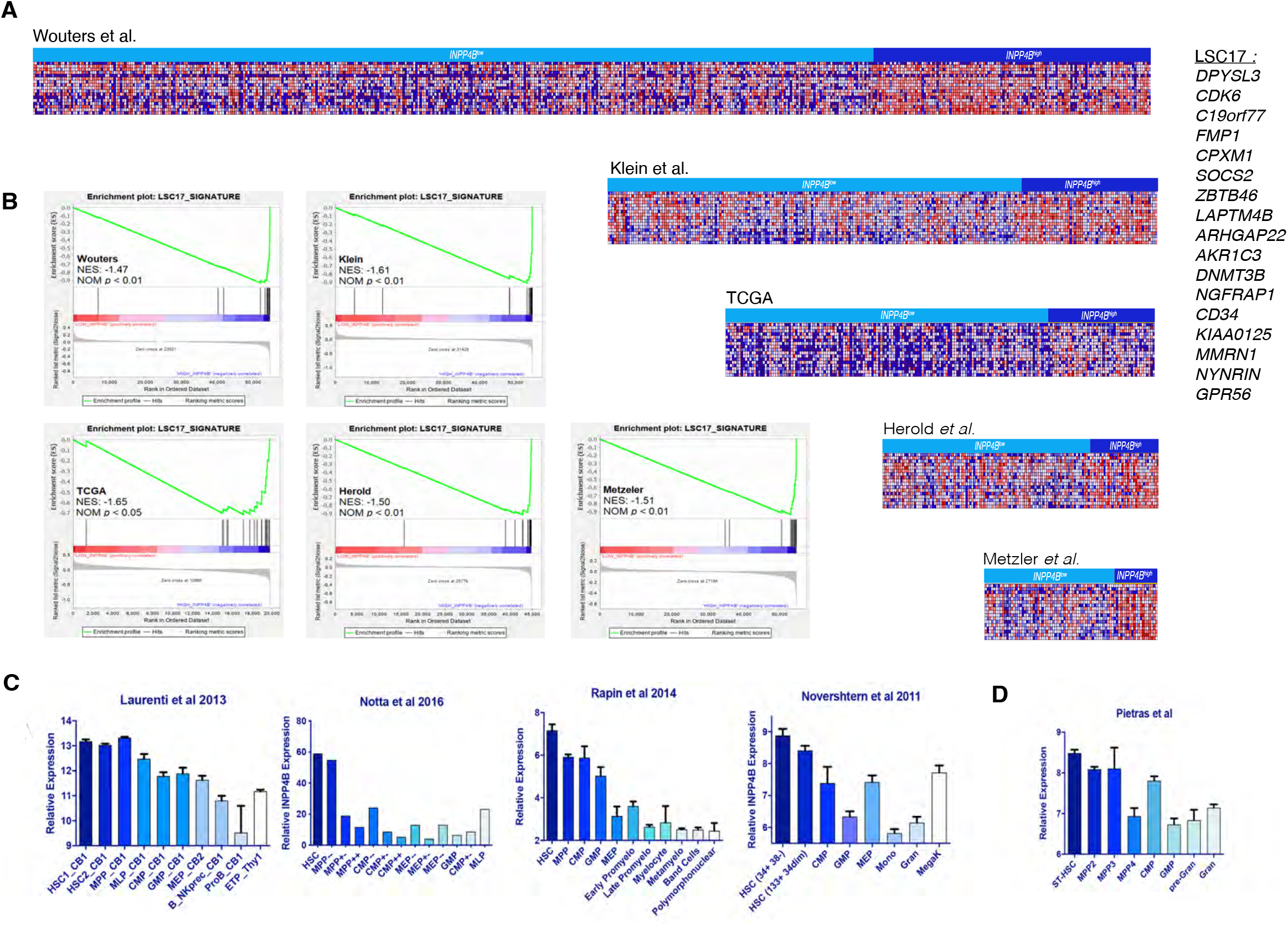
A. Heatmaps of LSC-17 gene expression in AML patient datasets, ranked by INPP4B expression (INPP4B-high quartile highlighted dark blue). B. Gene set enrichment analysis (GSEA) enrichment plot of the LSC-17 gene set in the Wouters, Klein, TCGA, Herold and Metzeler AML patient datasets ranked by INPP4B AML expression. C. Relative expression of INPP4B in sorted human or (D) sorted mouse hematopoietic stem and progenitor cells.

**Supplementary Figure 3.**
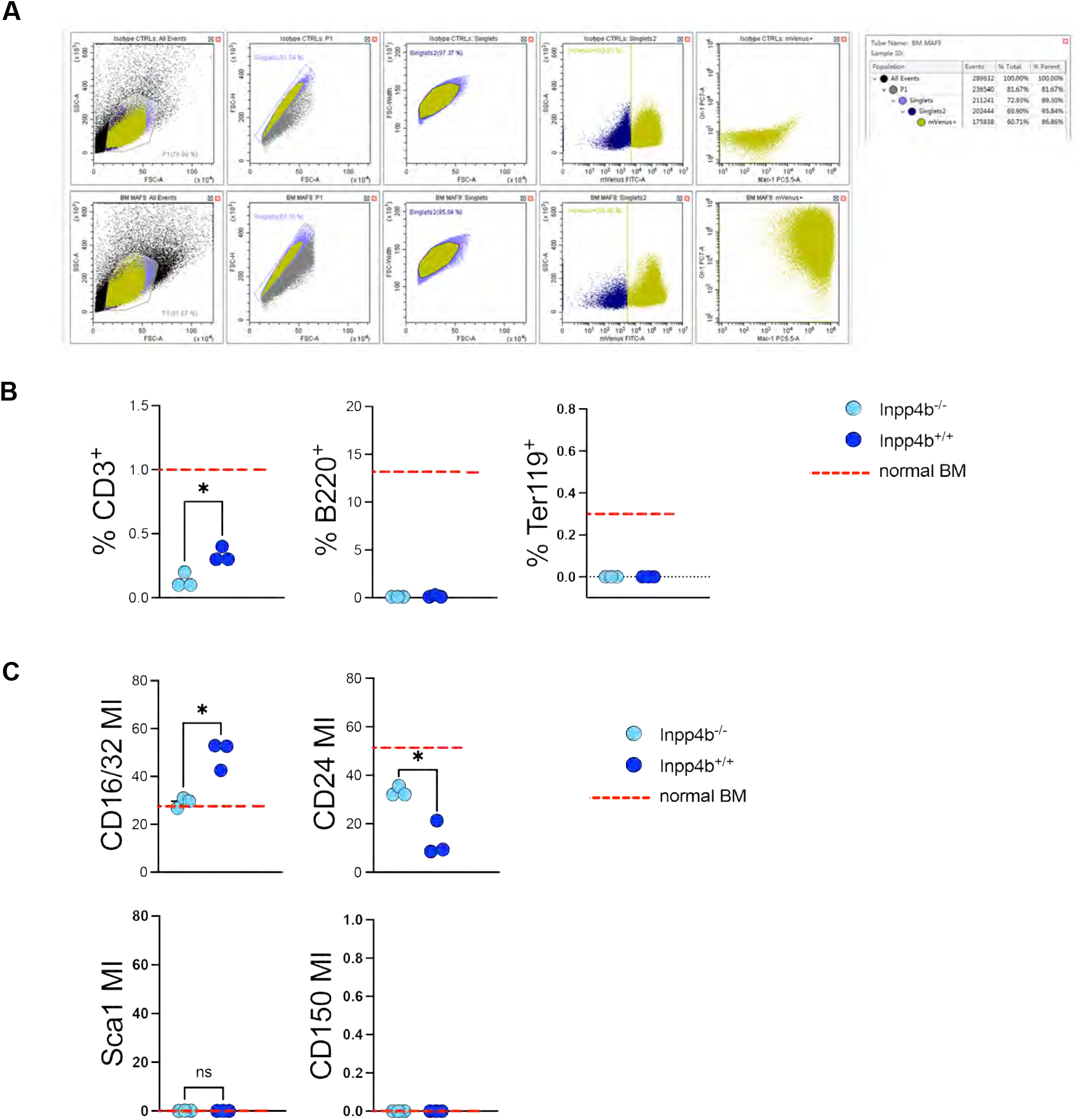
A. Gating strategy for generation of MLL-AF9 murine leukemias from age- and sex-matched adult Inpp4b+/+ and Inpp4b-/- mice, expressing both Gr-1 and Mac-1 surface markers. B. Scatter plots of % positive cell surface markers shown. T-test performed with Welch’s correction. C. Scatter plots of median intensity of cell surface markers shown. T-test performed with Welch’s correction.

**Supplementary Figure 4.**
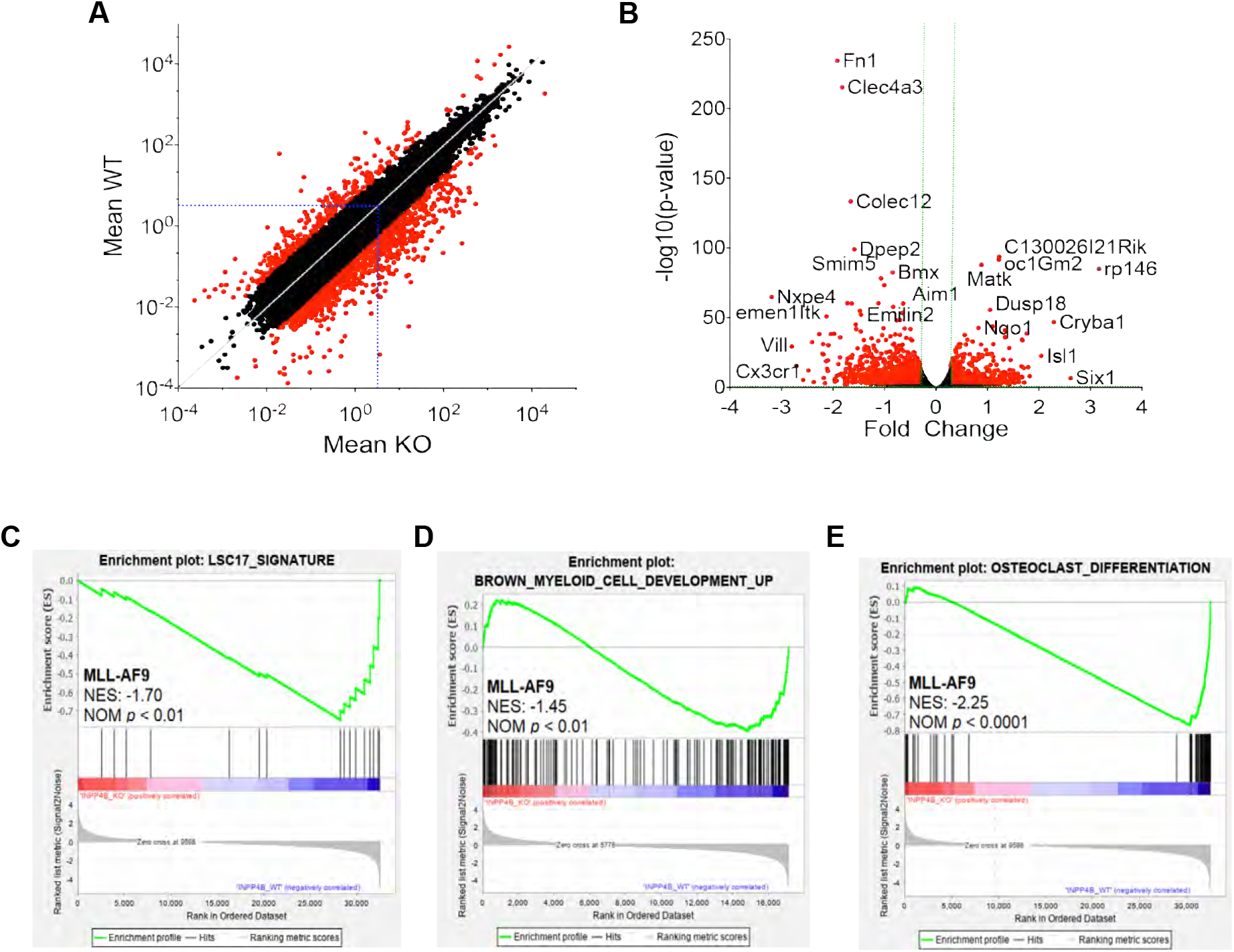
A. Scatter plot of TPM (transcript per million) values from genes with significantly altered expression in Inpp4b+/+ and Inpp4b-/- MLL-AF9 leukemia blast cells (n = 3 vs. 3, FDR < 0.01). B. Volcano plot of all genes from Inpp4b+/+ and Inpp4b-/- MLL-AF9 leukemia blast cells (n = 3 vs. 3, FDR < 0.01). GSEA enrichment plots demonstrating relationship between differentially expressed geneds in Inpp4b+/+ and Inpp4b-/- MLL-AF9 leukemia blast cells and (C) LSC17 signature, (D) KEGG myeloid cell development genes (E) and KEGG_osteoclast differentiation genes.

**Supplementary Figure 5.**
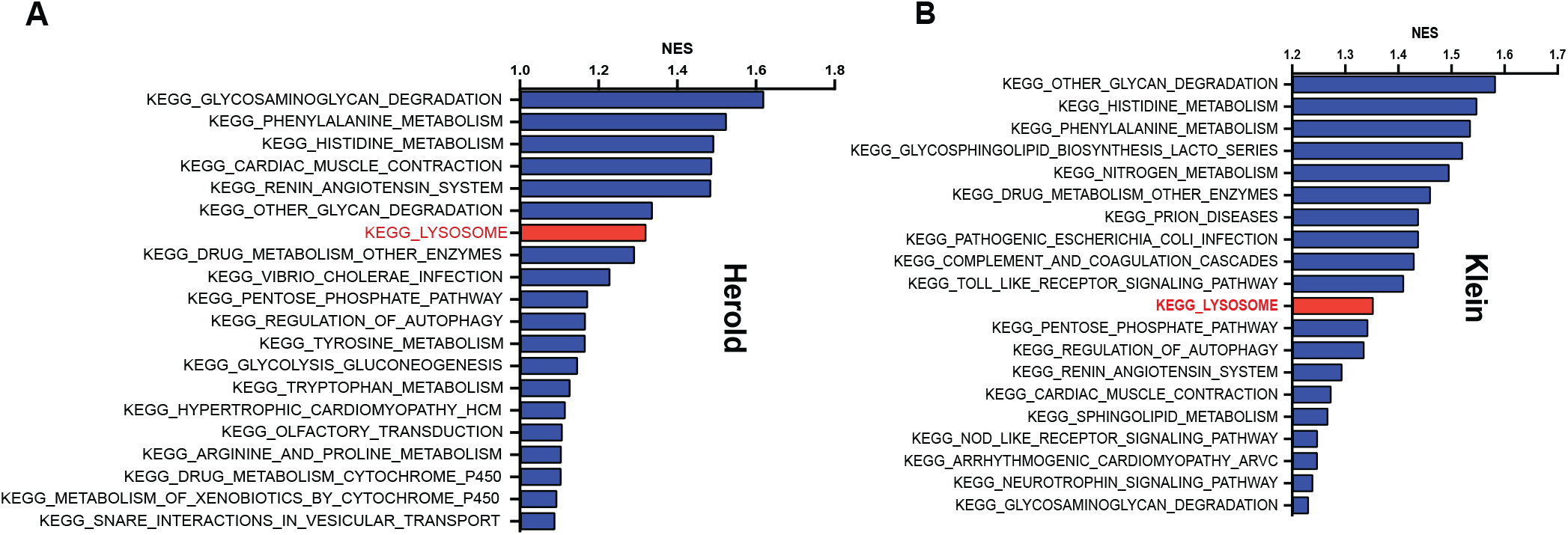
Histograms of the top 20 enriched KEGG gene sets in INPP4B-low patients from the (A) Herold, (B) Klein patient datasets ranked by normalized enrichment score (NES).

**Supplementary Figure 6.**
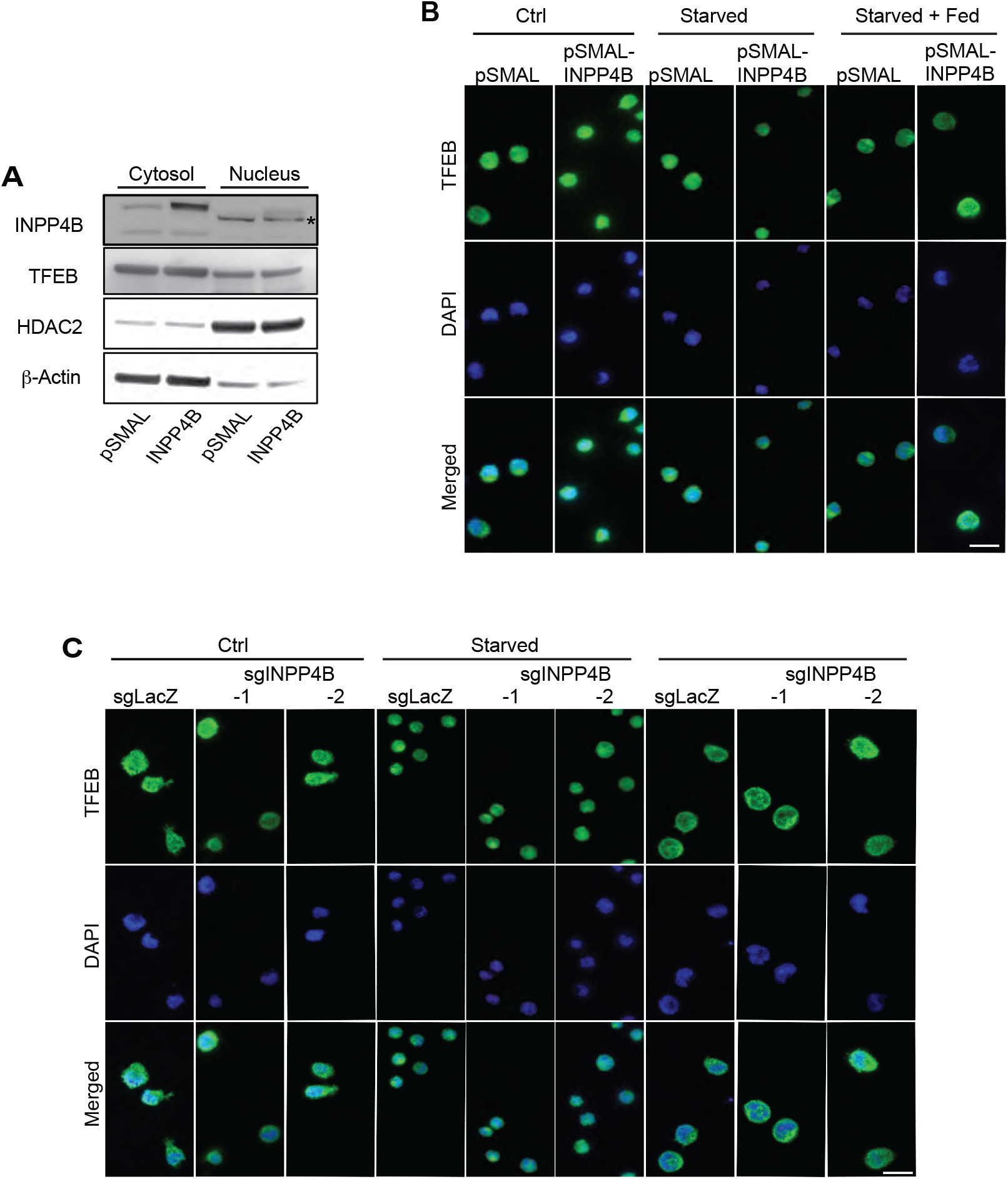
A. Western blot of OCI-AML2 cells’ cytosolic and nuclear extracts. B. OCI-AML2 INPP4B overexpressing and (C) knockout cells. Steady state control, FCS starved or starved and FCS fed cells were immunostained for TFEB and counterstained with DAPI.

**Supplementary Figure 7.**
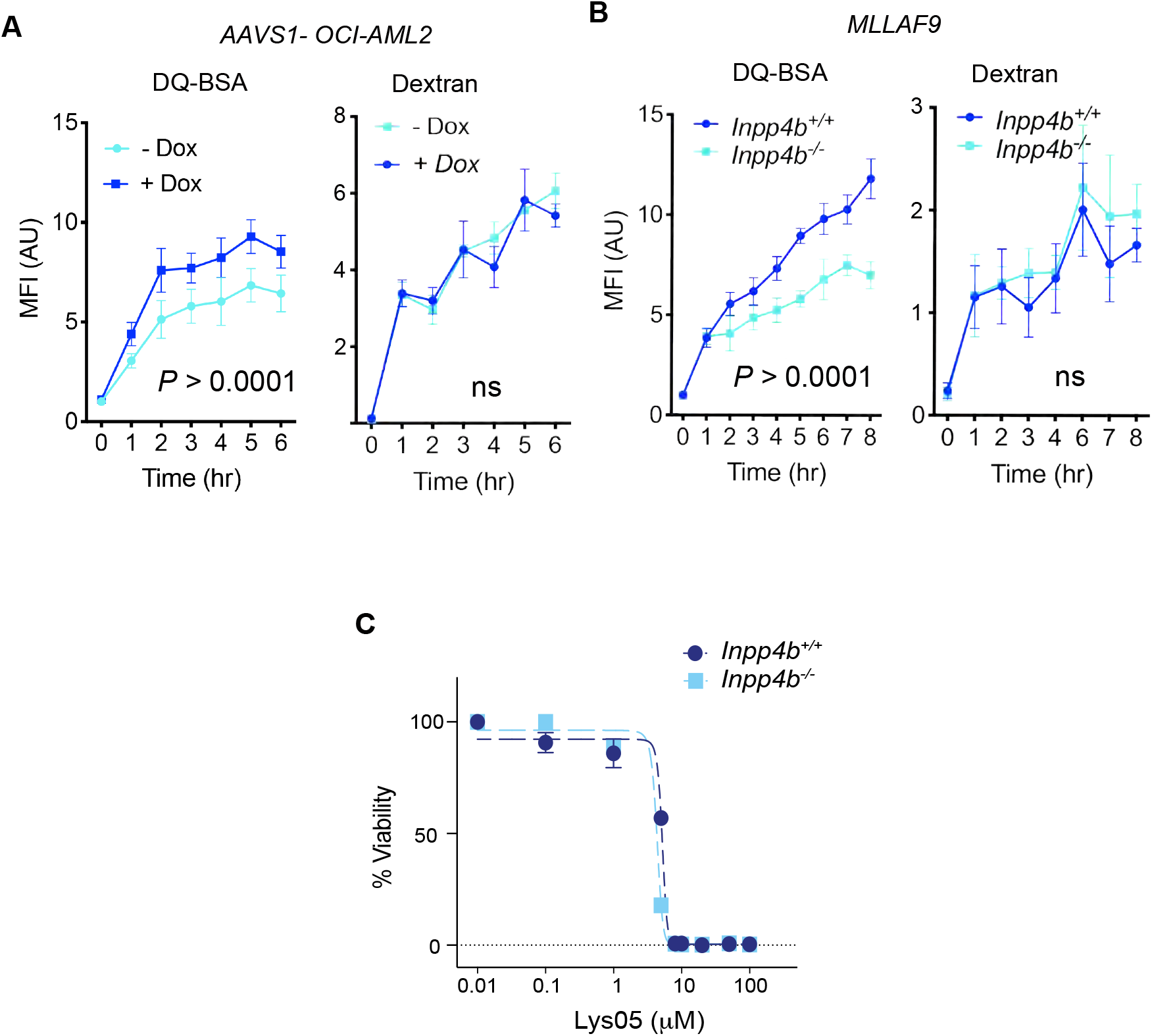
INPP4B regulates lysosomal biology in leukemia cells. A. Inducible-INPP4B OCI-AML2 cells were treated without or with doxycycline (100mg/mL) and incubated with DQ-BSA-Green™ or Dextran-Red. Fluorescence was monitored by flow cytometry hourly, for up to 6 hours. Full time course displayed. B. Inpp4b+/+ and Inpp4B-/- MLL-AF9 leukemia blasts were incubated with and incubated with DQ-BSA-Green™ or Dextran-Red. Fluorescence was monitored by flow cytometry hourly, for up to 8 hours. Full time course displayed. C. Lys05 Dose response in Inpp4b+/+ and Inpp4B-/- MLL-AF9 leukemia blasts.

**Supplementary Figure 8:**
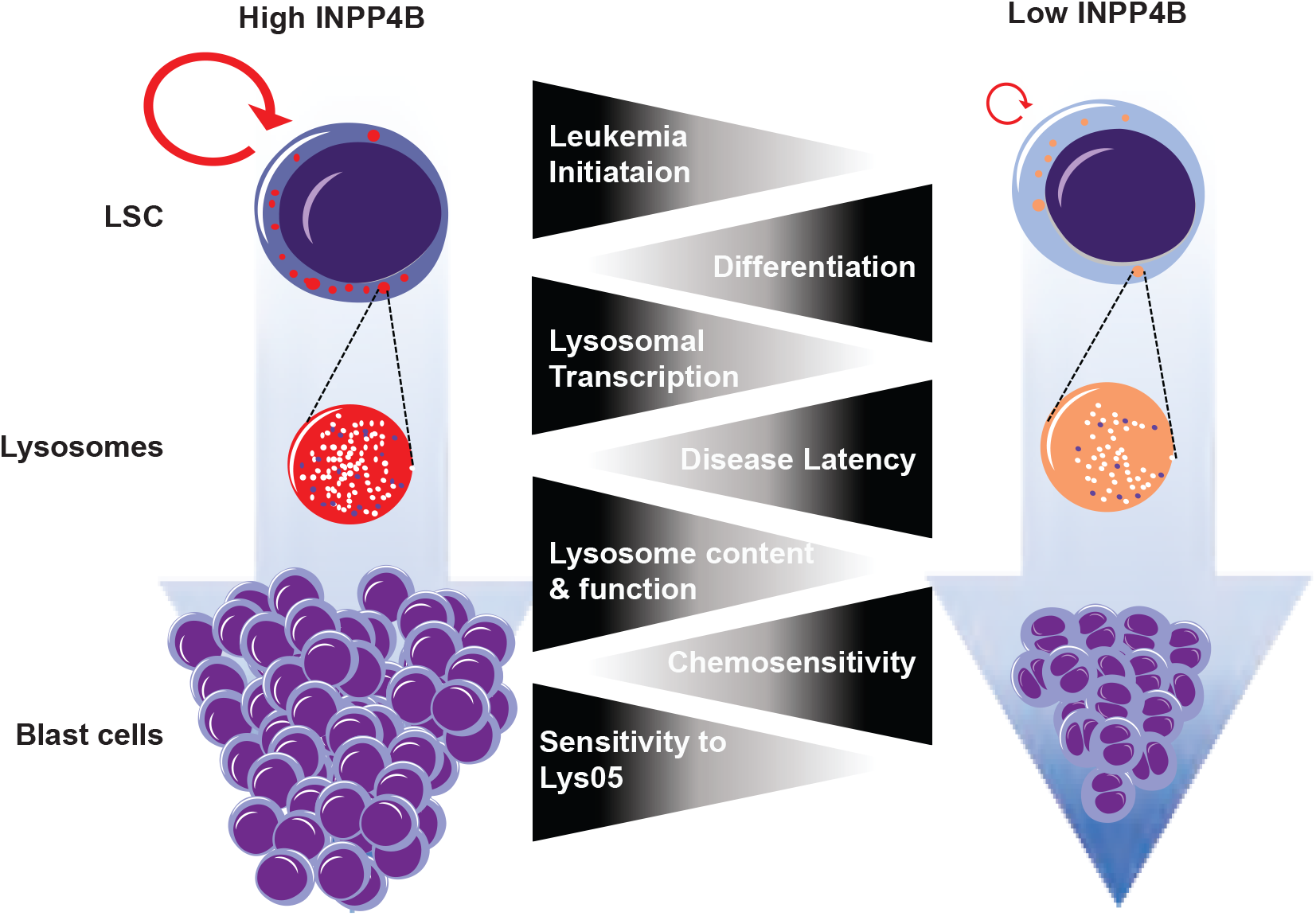
Model of INPP4B function in leukemia stem cells and leukemic blasts.

**SUPPLEMENTARY TABLE 1.**
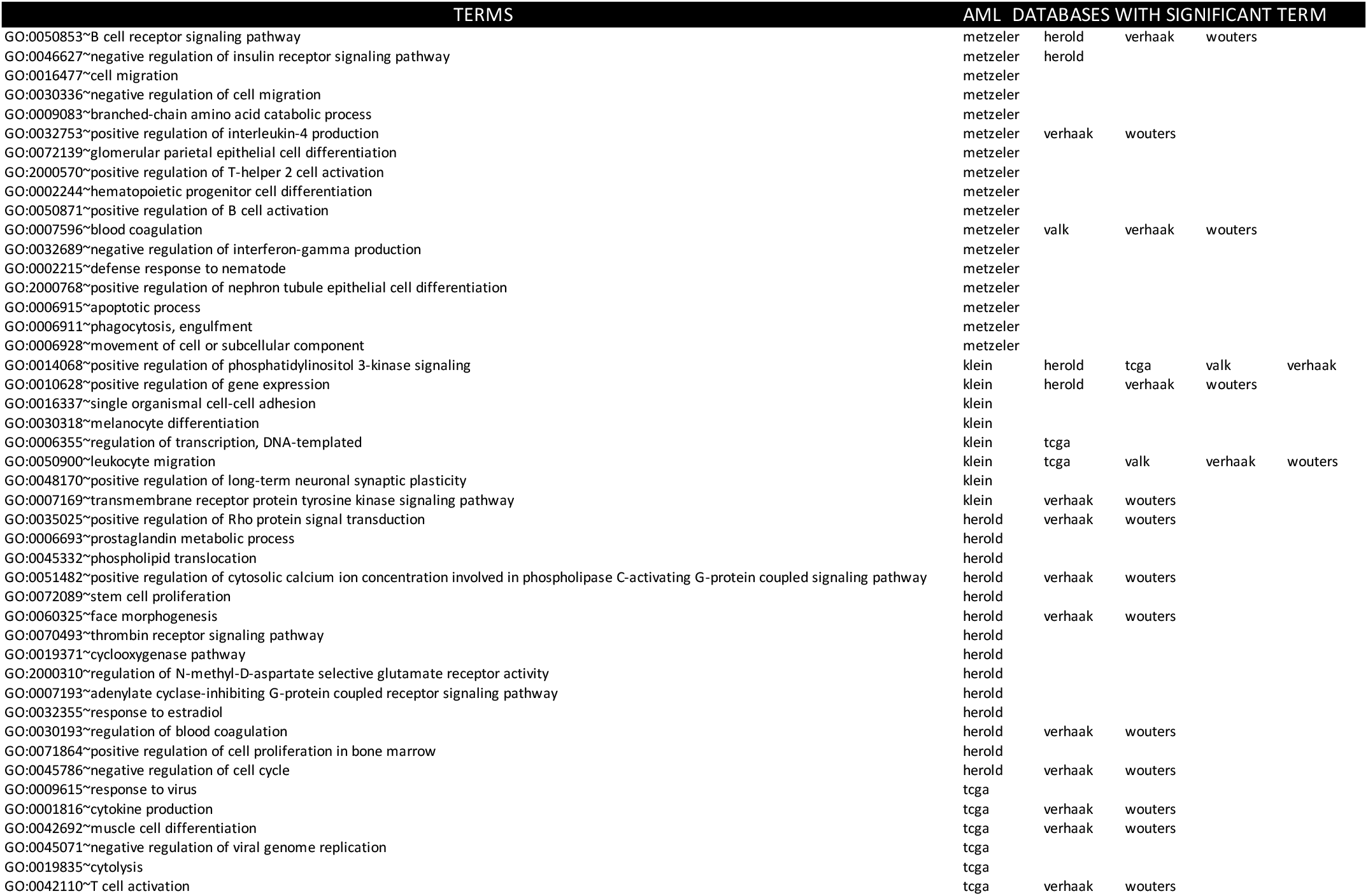
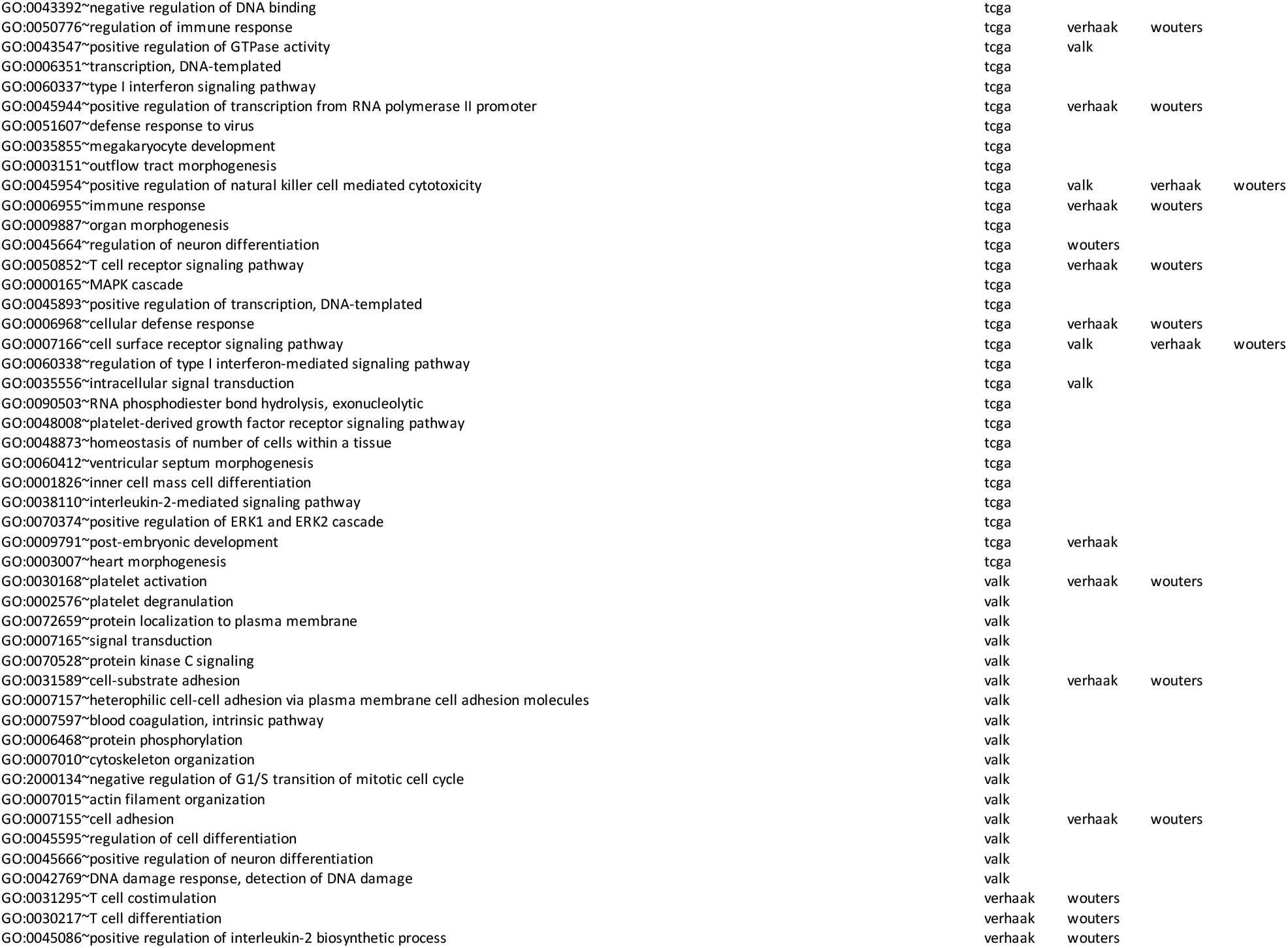
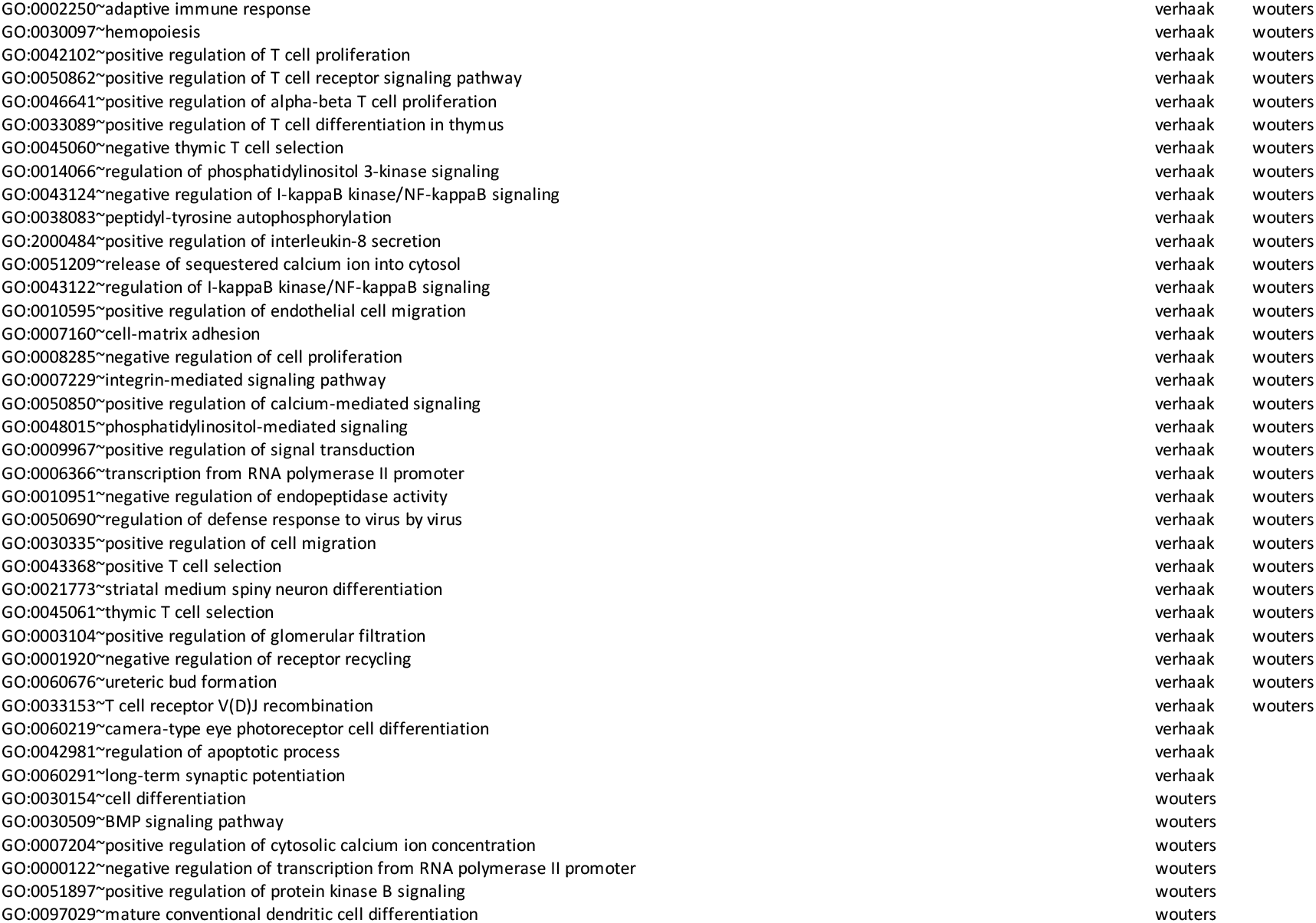

| TERMS | AML DATABASES WITH SIGNIFICANT TERM |  |  |  |  |
| --- | --- | --- | --- | --- | --- |
| GO:0050853~B cell receptor signaling pathway | metzeler | herold | verhaak | wouters |  |
| GO:0046627~negative regulation of insulin receptor signaling pathway | metzeler | herold |  |  |  |
| GO:0016477~cell migration | metzeler |  |  |  |  |
| GO:0030336~negative regulation of cell migration | metzeler |  |  |  |  |
| GO:0009083~branched-chain amino acid catabolic process | metzeler |  |  |  |  |
| GO:0032753~positive regulation of interleukin-4 production | metzeler | verhaak | wouters |  |  |
| GO:0072139~glomerular parietal epithelial cell differentiation | metzeler |  |  |  |  |
| GO:2000570~positive regulation of T-helper 2 cell activation | metzeler |  |  |  |  |
| GO:0002244~hematopoietic progenitor cell differentiation | metzeler |  |  |  |  |
| GO:0050871~positive regulation of B cell activation | metzeler |  |  |  |  |
| GO:0007596~blood coagulation | metzeler | valk | verhaak | wouters |  |
| GO:0032689~negative regulation of interferon-gamma production | metzeler |  |  |  |  |
| GO:0002215~defense response to nematode | metzeler |  |  |  |  |
| GO:2000768~positive regulation of nephron tubule epithelial cell differentiation | metzeler |  |  |  |  |
| GO:0006915~apoptotic process | metzeler |  |  |  |  |
| GO:0006911~phagocytosis, engulfment | metzeler |  |  |  |  |
| GO:0006928~movement of cell or subcellular component | metzeler |  |  |  |  |
| GO:0014068~positive regulation of phosphatidylinositol 3-kinase signaling | klein | herold | tcga | valk | verhaak |
| GO:0010628~positive regulation of gene expression | klein | herold | verhaak | wouters |  |
| GO:0016337~single organismal cell-cell adhesion | klein |  |  |  |  |
| GO:0030318~melanocyte differentiation | klein |  |  |  |  |
| GO:0006355~regulation of transcription, DNA-templated | klein | tcga |  |  |  |
| GO:0050900~leukocyte migration | klein | tcga | valk | verhaak | wouters |
| GO:0048170~positive regulation of long-term neuronal synaptic plasticity | klein |  |  |  |  |
| GO:0007169~transmembrane receptor protein tyrosine kinase signaling pathway | klein | verhaak | wouters |  |  |
| GO:0035025~positive regulation of Rho protein signal transduction | herold | verhaak | wouters |  |  |
| GO:0006693~prostaglandin metabolic process | herold |  |  |  |  |
| GO:0045332~phospholipid translocation | herold |  |  |  |  |
| GO:0051482~positive regulation of cytosolic calcium ion concentration involved in phospholipase C-activating G-protein coupled signaling pathway | herold | verhaak | wouters |  |  |
| GO:0072089~stem cell proliferation | herold |  |  |  |  |
| GO:0060325~face morphogenesis | herold | verhaak | wouters |  |  |
| GO:0070493~thrombin receptor signaling pathway | herold |  |  |  |  |
| GO:0019371~cyclooxygenase pathway | herold |  |  |  |  |
| GO:2000310~regulation of N-methyl-D-aspartate selective glutamate receptor activity | herold |  |  |  |  |
| GO:0007193~adenylate cyclase-inhibiting G-protein coupled receptor signaling pathway | herold |  |  |  |  |
| GO:0032355~response to estradiol | herold |  |  |  |  |
| GO:0030193~regulation of blood coagulation | herold | verhaak | wouters |  |  |
| GO:0071864~positive regulation of cell proliferation in bone marrow | herold |  |  |  |  |
| GO:0045786~negative regulation of cell cycle | herold | verhaak | wouters |  |  |
| GO:0009615~response to virus | tcga |  |  |  |  |
| GO:0001816~cytokine production | tcga | verhaak | wouters |  |  |
| GO:0042692~muscle cell differentiation | tcga | verhaak | wouters |  |  |
| GO:0045071~negative regulation of viral genome replication | tcga |  |  |  |  |
| GO:0019835~cytolysis | tcga |  |  |  |  |
| GO:0042110~T cell activation | tcga | verhaak | wouters |  |  |

|  |  |  |  |  |
| --- | --- | --- | --- | --- |
| GO:0043392~negative regulation of DNA binding | tcga |  |  |  |
| GO:0050776~regulation of immune response | tcga | verhaak | wouters |  |
| GO:0043547~positive regulation of GTPase activity | tcga | valk |  |  |
| GO:0006351~transcription, DNA-templated | tcga |  |  |  |
| GO:0060337~type I interferon signaling pathway | tcga |  |  |  |
| GO:0045944~positive regulation of transcription from RNA polymerase II promoter | tcga | verhaak | wouters |  |
| GO:0051607~defense response to virus | tcga |  |  |  |
| GO:0035855~megakaryocyte development | tcga |  |  |  |
| GO:0003151~outflow tract morphogenesis | tcga |  |  |  |
| GO:0045954~positive regulation of natural killer cell mediated cytotoxicity | tcga | valk | verhaak | wouters |
| GO:0006955~immune response | tcga | verhaak | wouters |  |
| GO:0009887~organ morphogenesis | tcga |  |  |  |
| GO:0045664~regulation of neuron differentiation | tcga | wouters |  |  |
| GO:0050852~T cell receptor signaling pathway | tcga | verhaak | wouters |  |
| GO:0000165~MAPK cascade | tcga |  |  |  |
| GO:0045893~positive regulation of transcription, DNA-templated | tcga |  |  |  |
| GO:0006968~cellular defense response | tcga | verhaak | wouters |  |
| GO:0007166~cell surface receptor signaling pathway | tcga | valk | verhaak | wouters |
| GO:0060338~regulation of type I interferon-mediated signaling pathway | tcga |  |  |  |
| GO:0035556~intracellular signal transduction | tcga | valk |  |  |
| GO:0090503~RNA phosphodiester bond hydrolysis, exonucleolytic | tcga |  |  |  |
| GO:0048008~platelet-derived growth factor receptor signaling pathway | tcga |  |  |  |
| GO:0048873~homeostasis of number of cells within a tissue | tcga |  |  |  |
| GO:0060412~ventricular septum morphogenesis | tcga |  |  |  |
| GO:0001826~inner cell mass cell differentiation | tcga |  |  |  |
| GO:0038110~interleukin-2-mediated signaling pathway | tcga |  |  |  |
| GO:0070374~positive regulation of ERK1 and ERK2 cascade | tcga |  |  |  |
| GO:0009791~post-embryonic development | tcga | verhaak |  |  |
| GO:0003007~heart morphogenesis | tcga |  |  |  |
| GO:0030168~platelet activation | valk | verhaak | wouters |  |
| GO:0002576~platelet degranulation | valk |  |  |  |
| GO:0072659~protein localization to plasma membrane | valk |  |  |  |
| GO:0007165~signal transduction | valk |  |  |  |
| GO:0070528~protein kinase C signaling | valk |  |  |  |
| GO:0031589~cell-substrate adhesion | valk | verhaak | wouters |  |
| GO:0007157~heterophilic cell-cell adhesion via plasma membrane cell adhesion molecules | valk |  |  |  |
| GO:0007597~blood coagulation, intrinsic pathway | valk |  |  |  |
| GO:0006468~protein phosphorylation | valk |  |  |  |
| GO:0007010~cytoskeleton organization | valk |  |  |  |
| GO:2000134~negative regulation of G1/S transition of mitotic cell cycle | valk |  |  |  |
| GO:0007015~actin filament organization | valk |  |  |  |
| GO:0007155~cell adhesion | valk | verhaak | wouters |  |
| GO:0045595~regulation of cell differentiation | valk |  |  |  |
| GO:0045666~positive regulation of neuron differentiation | valk |  |  |  |
| GO:0042769~DNA damage response, detection of DNA damage | valk |  |  |  |
| GO:0031295~T cell costimulation | verhaak | wouters |  |  |
| GO:0030217~T cell differentiation | verhaak | wouters |  |  |
| GO:0045086~positive regulation of interleukin-2 biosynthetic process | verhaak | wouters |  |  |

|  |  |  |
| --- | --- | --- |
| GO:0002250~adaptive immune response | verhaak | wouters |
| GO:0030097~hemopoiesis | verhaak | wouters |
| GO:0042102~positive regulation of T cell proliferation | verhaak | wouters |
| GO:0050862~positive regulation of T cell receptor signaling pathway | verhaak | wouters |
| GO:0046641~positive regulation of alpha-beta T cell proliferation | verhaak | wouters |
| GO:0033089~positive regulation of T cell differentiation in thymus | verhaak | wouters |
| GO:0045060~negative thymic T cell selection | verhaak | wouters |
| GO:0014066~regulation of phosphatidylinositol 3-kinase signaling | verhaak | wouters |
| GO:0043124~negative regulation of I-kappaB kinase/NF-kappaB signaling | verhaak | wouters |
| GO:0038083~peptidyl-tyrosine autophosphorylation | verhaak | wouters |
| GO:2000484~positive regulation of interleukin-8 secretion | verhaak | wouters |
| GO:0051209~release of sequestered calcium ion into cytosol | verhaak | wouters |
| GO:0043122~regulation of I-kappaB kinase/NF-kappaB signaling | verhaak | wouters |
| GO:0010595~positive regulation of endothelial cell migration | verhaak | wouters |
| GO:0007160~cell-matrix adhesion | verhaak | wouters |
| GO:0008285~negative regulation of cell proliferation | verhaak | wouters |
| GO:0007229~integrin-mediated signaling pathway | verhaak | wouters |
| GO:0050850~positive regulation of calcium-mediated signaling | verhaak | wouters |
| GO:0048015~phosphatidylinositol-mediated signaling | verhaak | wouters |
| GO:0009967~positive regulation of signal transduction | verhaak | wouters |
| GO:0006366~transcription from RNA polymerase II promoter | verhaak | wouters |
| GO:0010951~negative regulation of endopeptidase activity | verhaak | wouters |
| GO:0050690~regulation of defense response to virus by virus | verhaak | wouters |
| GO:0030335~positive regulation of cell migration | verhaak | wouters |
| GO:0043368~positive T cell selection | verhaak | wouters |
| GO:0021773~striatal medium spiny neuron differentiation | verhaak | wouters |
| GO:0045061~thymic T cell selection | verhaak | wouters |
| GO:0003104~positive regulation of glomerular filtration | verhaak | wouters |
| GO:0001920~negative regulation of receptor recycling | verhaak | wouters |
| GO:0060676~ureteric bud formation | verhaak | wouters |
| GO:0033153~T cell receptor V(D)J recombination | verhaak | wouters |
| GO:0060219~camera-type eye photoreceptor cell differentiation | verhaak |  |
| GO:0042981~regulation of apoptotic process | verhaak |  |
| GO:0060291~long-term synaptic potentiation | verhaak |  |
| GO:0030154~cell differentiation | wouters |  |
| GO:0030509~BMP signaling pathway | wouters |  |
| GO:0007204~positive regulation of cytosolic calcium ion concentration | wouters |  |
| GO:0000122~negative regulation of transcription from RNA polymerase II promoter | wouters |  |
| GO:0051897~positive regulation of protein kinase B signaling | wouters |  |
| GO:0097029~mature conventional dendritic cell differentiation | wouters |  |

**Supplementary Table 2.**
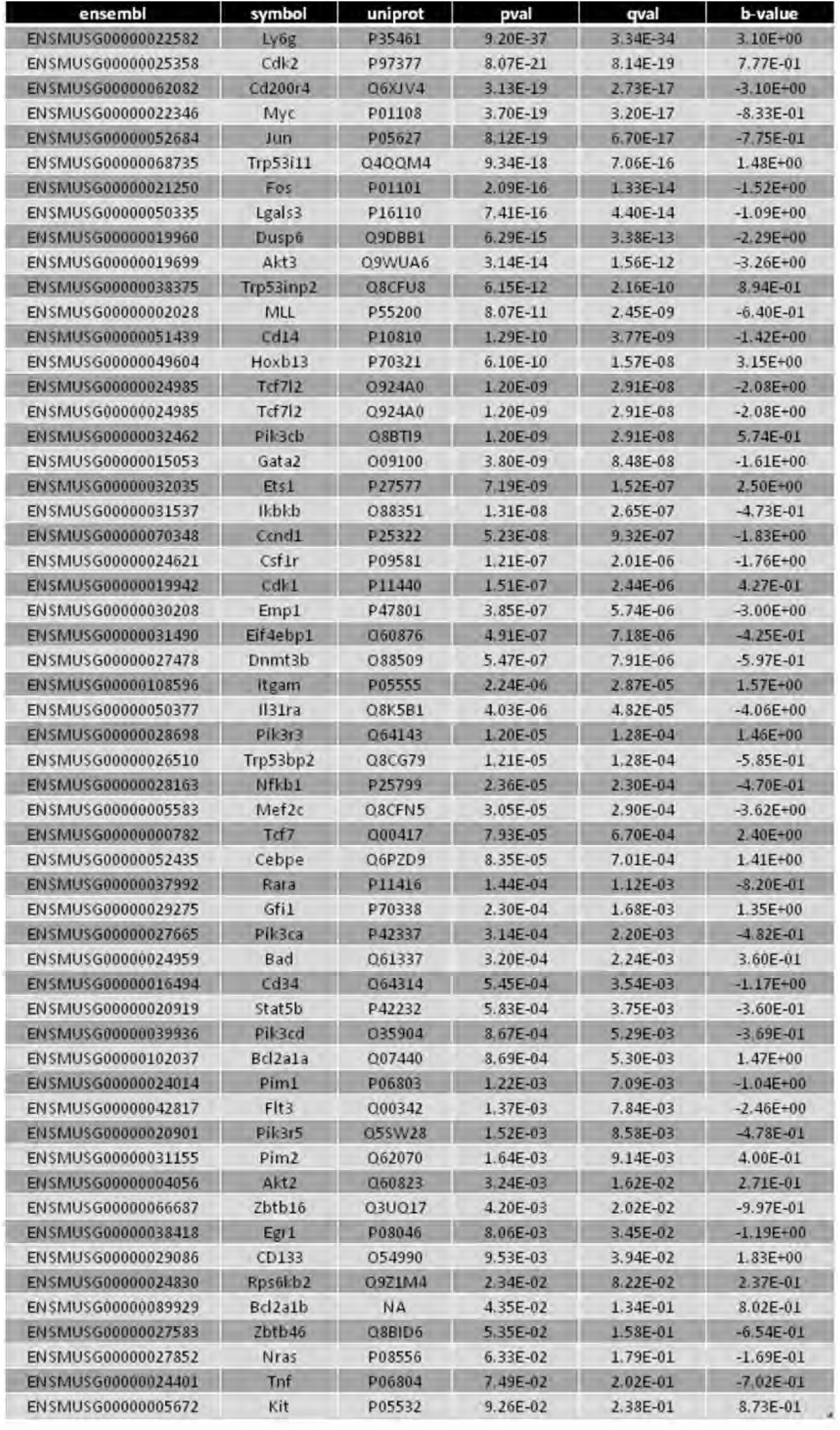

**Supplementary Table 3.**

Woolley et al. Supplementary Table 3
| Name | Id | Source Name | Organism | Uploaded gene count | Genes in InnateDB for this entity | P-value | P-value (corrected) |
| --- | --- | --- | --- | --- | --- | --- | --- |
| Cell cycle | 122 | KEGG | 10090 | 72 | 123 | 7.60E-13 | 2.03E-10 |
| **DNA replication** | **95** | **KEGG** | **10090** | **28** | **35** | **1.96E-10** | **3.94E-08** |
| Lysosome | 132 | KEGG | 10090 | 63 | 124 | 4.50E-08 | 3.62E-06 |
| **Homologous recombination** | **103** | **KEGG** | **10090** | **27** | **41** | **4.30E-07** | **3.00E-05** |
| Osteoclast differentiation | 156 | KEGG | 10090 | 61 | 125 | 4.60E-07 | 3.08E-05 |
| **NOD-like receptor signaling pathway** | **171** | **KEGG** | **10090** | **73** | **166** | **5.31E-06** | **2.75E-04** |
| Cellular senescence | 145 | KEGG | 10090 | 75 | 177 | 1.98E-05 | 8.61E-04 |
| **TNF signaling pathway** | **185** | **KEGG** | **10090** | **50** | **107** | **2.22E-05** | **9.38E-04** |
| MicroRNAs in cancer | 283 | KEGG | 10090 | 62 | 142 | 3.45E-05 | 0.001386308 |
| **Toll-like receptor signaling pathway** | **170** | **KEGG** | **10090** | **46** | **98** | **4.06E-05** | **0.00158865** |
| Oocyte meiosis | 123 | KEGG | 10090 | 52 | 115 | 4.69E-05 | 0.001750363 |
| **HTLV-I infection** | **274** | **KEGG** | **10090** | **106** | **274** | **5.30E-05** | **0.001892981** |
| Apoptosis | 139 | KEGG | 10090 | 59 | 136 | 6.61E-05 | 0.002309443 |
| **Nucleotide excision repair** | **101** | **KEGG** | **10090** | **24** | **43** | **1.03E-04** | **0.003172607** |
| Fluid shear stress and atherosclerosis | 315 | KEGG | 10090 | 60 | 142 | 1.42E-04 | 0.003867482 |
| **Base excision repair** | **100** | **KEGG** | **10090** | **20** | **34** | **1.46E-04** | **0.003909688** |
| Platelet activation | 167 | KEGG | 10090 | 53 | 123 | 1.88E-04 | 0.004781539 |
| **Fanconi anemia pathway** | **105** | **KEGG** | **10090** | **27** | **52** | **2.00E-04** | **0.005014235** |
| Mismatch repair | 102 | KEGG | 10090 | 15 | 23 | 2.03E-04 | 0.004941126 |

## References

Amaravadi, R.K., and Winkler, J.D. (2012). Lys05: a new lysosomal autophagy inhibitor. Autophagy 8, 1383–1384.

Balgobind, B.V., Van den Heuvel-Eibrink, M.M., De Menezes, R.X., Reinhardt, D., Hollink, I.H.I.M., Arentsen-Peters, S.T.J.C.M., van Wering, E.R., Kaspers, G.J.L., Cloos, J., de Bont, E.S.J.M., et al. (2011). Evaluation of gene expression signatures predictive of cytogenetic and molecular subtypes of pediatric acute myeloid leukemia. Haematologica 96, 221–230.

Bernard, D., Gebbia, M., Prabha, S., Gronda, M., MacLean, N., Wang, X., Hurren, R., Sukhai, M.A., Cho, E.E., Manolson, M.F., et al. (2015). Select microtubule inhibitors increase lysosome acidity and promote lysosomal disruption in acute myeloid leukemia (AML) cells. Apoptosis 20, 948–959.

Cancer Genome Atlas Research Network, Ley, T.J., Miller, C., Ding, L., Raphael, B.J., Mungall, A.J., Robertson, A.G., Hoadley, K., Triche, T.J., Laird, P.W., et al. (2013). Genomic and epigenomic landscapes of adult de novo acute myeloid leukemia. N. Engl. J. Med. 368, 2059–2074.

Cechakova, L., Ondrej, M., Pavlik, V., Jost, P., Cizkova, D., Bezrouk, A., Pejchal, J., Amaravadi, R.K., Winkler, J.D., and Tichy, A. (2019). A potent autophagy inhibitor (lys05) enhances the impact of ionizing radiation on human lung cancer cells H1299. Int. J. Mol. Sci. 20.

Chi, M.N., Guo, S.T., Wilmott, J.S., Guo, X.Y., Yan, X.G., Wang, C.Y., Liu, X.Y., Jin, L., Tseng, H.-Y., Liu, T., et al. (2015). INPP4B is upregulated and functions as an oncogenic driver through SGK3 in a subset of melanomas. Oncotarget 6, 39891–39907.

Dick, J.E. (2008). Stem cell concepts renew cancer research. Blood 112, 4793–4807.

Dzneladze, I., He, R., Woolley, J.F., Son, M.H., Sharobim, M.H., Greenberg, S.A., Gabra, M., Langlois, C., Rashid, A., Hakem, A., et al. (2015). INPP4B overexpression is associated with poor clinical outcome and therapy resistance in acute myeloid leukemia. Leukemia 29, 1485–1495.

Dzneladze, I., Woolley, J.F., Rossell, C., Han, Y., Rashid, A., Jain, M., Reimand, J., Minden, M.D., and Salmena, L. (2018). SubID, a non-median dichotomization tool for heterogeneous populations, reveals the pan-cancer significance of INPP4B and its regulation by EVI1 in AML. PLoS One 13, e0191510.

Eppert, K., Takenaka, K., Lechman, E.R., Waldron, L., Nilsson, B., van Galen, P., Metzeler, K.H., Poeppl, A., Ling, V., Beyene, J., et al. (2011). Stem cell gene expression programs influence clinical outcome in human leukemia. Nat. Med. 17, 1086–1093.

Ferron, M., Boudiffa, M., Arsenault, M., Rached, M., Pata, M., Giroux, S., Elfassihi, L., Kisseleva, M., Majerus, P.W., Rousseau, F., et al. (2011). Inositol polyphosphate 4-phosphatase B as a regulator of bone mass in mice and humans. Cell Metab. 14, 466–477.

Frenquelli, M., and Tonon, G. (2020). WNT signaling in hematological malignancies. Front. Oncol. 10, 615190.

García-Prat, L., Kaufmann, K.B., Schneiter, F., Voisin, V., Murison, A., Chen, J., Chan-Seng-Yue, M., Gan, O.I., McLeod, J.L., Smith, S.A., et al. (2021). TFEB-mediated endolysosomal activity controls human hematopoietic stem cell fate. Cell Stem Cell 28, 1838–1850.e10.

Gasser, J.A., Inuzuka, H., Lau, A.W., Wei, W., Beroukhim, R., and Toker, A. (2014). SGK3 mediates INPP4B-dependent PI3K signaling in breast cancer. Mol. Cell 56, 595–607.

Grainger, S., Traver, D., and Willert, K. (2018). Wnt signaling in hematological malignancies. Prog Mol Biol Transl Sci 153, 321–341.

Gu, Z., Gu, L., Eils, R., Schlesner, M., and Brors, B. (2014). circlize Implements and enhances circular visualization in R. Bioinformatics 30, 2811–2812.

Guo, S.T., Chi, M.N., Yang, R.H., Guo, X.Y., Zan, L.K., Wang, C.Y., Xi, Y.F., Jin, L., Croft, A., Tseng, H.Y., et al. (2016). INPP4B is an oncogenic regulator in human colon cancer. Oncogene 35, 3049–3061.

Hanahan, D. (2022). Hallmarks of cancer: new dimensions. Cancer Discov. 12, 31–46.

Herold, T., Jurinovic, V., Batcha, A.M.N., Bamopoulos, S.A., Rothenberg-Thurley, M., Ksienzyk, B., Hartmann, L., Greif, P.A., Phillippou-Massier, J., Krebs, S., et al. (2018). A 29-gene and cytogenetic score for the prediction of resistance to induction treatment in acute myeloid leukemia. Haematologica 103, 456–465.

Hu, Y., and Smyth, G.K. (2009). ELDA: extreme limiting dilution analysis for comparing depleted and enriched populations in stem cell and other assays. J. Immunol. Methods 347, 70–78.

Huang, D.W., Sherman, B.T., and Lempicki, R.A. (2009a). Systematic and integrative analysis of large gene lists using DAVID bioinformatics resources. Nat. Protoc. 4, 44–57.

Huang, D.W., Sherman, B.T., and Lempicki, R.A. (2009b). Bioinformatics enrichment tools: paths toward the comprehensive functional analysis of large gene lists. Nucleic Acids Res. 37, 1–13.

Inpanathan, S., and Botelho, R.J. (2019). The lysosome signaling platform: adapting with the times. Front. Cell Dev. Biol. 7, 113.

Jacomy, M., Venturini, T., Heymann, S., and Bastian, M. (2014). ForceAtlas2, a continuous graph layout algorithm for handy network visualization designed for the Gephi software. PLoS One 9, e98679.

Klein, H.-U., Ruckert, C., Kohlmann, A., Bullinger, L., Thiede, C., Haferlach, T., and Dugas, M. (2009). Quantitative comparison of microarray experiments with published leukemia related gene expression signatures. BMC Bioinformatics 10, 422.

Kreso, A., and Dick, J.E. (2014). Evolution of the cancer stem cell model. Cell Stem Cell 14, 275–291.

Lamming, D.W., and Bar-Peled, L. (2019). Lysosome: The metabolic signaling hub. Traffic 20, 27–38.

Lawrence, R.E., and Zoncu, R. (2019). The lysosome as a cellular centre for signalling, metabolism and quality control. Nat. Cell Biol. 21, 133–142.

Leeman, D.S., Hebestreit, K., Ruetz, T., Webb, A.E., McKay, A., Pollina, E.A., Dulken, B.W., Zhao, X., Yeo, R.W., Ho, T.T., et al. (2018). Lysosome activation clears aggregates and enhances quiescent neural stem cell activation during aging. Science 359, 1277–1283.

Loeffler, D., Wehling, A., Schneiter, F., Zhang, Y., Müller-Bötticher, N., Hoppe, P.S., Hilsenbeck, O., Kokkaliaris, K.D., Endele, M., and Schroeder, T. (2019). Publisher Correction: Asymmetric lysosome inheritance predicts activation of haematopoietic stem cells. Nature 573, E5.

Luzio, J.P., Rous, B.A., Bright, N.A., Pryor, P.R., Mullock, B.M., and Piper, R.C. (2000). Lysosome-endosome fusion and lysosome biogenesis. J. Cell Sci. 113 (Pt 9), 1515–1524.

Marwaha, R., and Sharma, M. (2017). DQ-Red BSA Trafficking Assay in Cultured Cells to Assess Cargo Delivery to Lysosomes. Bio Protoc 7.

McAfee, Q., Zhang, Z., Samanta, A., Levi, S.M., Ma, X.-H., Piao, S., Lynch, J.P., Uehara, T., Sepulveda, A.R., Davis, L.E., et al. (2012). Autophagy inhibitor Lys05 has single-agent antitumor activity and reproduces the phenotype of a genetic autophagy deficiency. Proc. Natl. Acad. Sci. USA 109, 8253–8258.

Metzeler, K.H., Hummel, M., Bloomfield, C.D., Spiekermann, K., Braess, J., Sauerland, M.-C., Heinecke, A., Radmacher, M., Marcucci, G., Whitman, S.P., et al. (2008). An 86-probe-set geneexpression signature predicts survival in cytogenetically normal acute myeloid leukemia. Blood 112, 4193–4201.

Ng, S.W.K., Mitchell, A., Kennedy, J.A., Chen, W.C., McLeod, J., Ibrahimova, N., Arruda, A., Popescu, A., Gupta, V., Schimmer, A.D., et al. (2016). A 17-gene stemness score for rapid determination of risk in acute leukaemia. Nature 540, 433–437.

Palmieri, M., Impey, S., Kang, H., di Ronza, A., Pelz, C., Sardiello, M., and Ballabio, A. (2011). Characterization of the CLEAR network reveals an integrated control of cellular clearance pathways. Hum. Mol. Genet. 20, 3852–3866.

Papaemmanuil, E., Gerstung, M., Bullinger, L., Gaidzik, V.I., Paschka, P., Roberts, N.D., Potter, N.E., Heuser, M., Thol, F., Bolli, N., et al. (2016). Genomic classification and prognosis in acute myeloid leukemia. N. Engl. J. Med. 374, 2209–2221.

Rafiq, S., McKenna, S.L., Muller, S., Tschan, M.P., and Humbert, M. (2021). Lysosomes in acute myeloid leukemia: potential therapeutic targets? Leukemia 35, 2759–2770.

Recher, C. (2015). INPP4B, a new player in the chemoresistance of AML. Blood 125, 2738–2739.

Rijal, S., Fleming, S., Cummings, N., Rynkiewicz, N.K., Ooms, L.M., Nguyen, N.-Y.N., Teh, T.-C., Avery, S., McManus, J.F., Papenfuss, A.T., et al. (2015). Inositol polyphosphate 4-phosphatase II (INPP4B) is associated with chemoresistance and poor outcome in AML. Blood 125, 2815–2824.

Rodgers, S.J., Ooms, L.M., Oorschot, V.M.J., Schittenhelm, R.B., Nguyen, E.V., Hamila, S.A., Rynkiewicz, N., Gurung, R., Eramo, M.J., Sriratana, A., et al. (2021). INPP4B promotes PI3Kα-dependent late endosome formation and Wnt/β-catenin signaling in breast cancer. Nat. Commun. 12, 3140.

Rouillard, A.D., Gundersen, G.W., Fernandez, N.F., Wang, Z., Monteiro, C.D., McDermott, M.G., and Ma’ayan, A. (2016). The harmonizome: a collection of processed datasets gathered to serve and mine knowledge about genes and proteins. Database (Oxford) 2016.

Sardiello, M., Palmieri, M., di Ronza, A., Medina, D.L., Valenza, M., Gennarino, V.A., Di Malta, C., Donaudy, F., Embrione, V., Polishchuk, R.S., et al. (2009). A gene network regulating lysosomal biogenesis and function. Science 325, 473–477.

Savini, M., Zhao, Q., and Wang, M.C. (2019). Lysosomes: signaling hubs for metabolic sensing and longevity. Trends Cell Biol. 29, 876–887.

Settembre, C., and Ballabio, A. (2014). Lysosome: regulator of lipid degradation pathways. Trends Cell Biol. 24, 743–750.

Siegel, R.L., Miller, K.D., and Jemal, A. (2018). Cancer statistics, 2018. CA Cancer J Clin 68, 7–30.

Somervaille, T.C.P., and Cleary, M.L. (2006). Identification and characterization of leukemia stem cells in murine MLL-AF9 acute myeloid leukemia. Cancer Cell 10, 257–268.

Sukhai, M.A., Prabha, S., Hurren, R., Rutledge, A.C., Lee, A.Y., Sriskanthadevan, S., Sun, H., Wang, X., Skrtic, M., Seneviratne, A., et al. (2013). Lysosomal disruption preferentially targets acute myeloid leukemia cells and progenitors. J. Clin. Invest. 123, 315–328.

Szklarczyk, D., Gable, A.L., Lyon, D., Junge, A., Wyder, S., Huerta-Cepas, J., Simonovic, M., Doncheva, N.T., Morris, J.H., Bork, P., et al. (2019). STRING v11: protein-protein association networks with increased coverage, supporting functional discovery in genome-wide experimental datasets. Nucleic Acids Res. 47, D607–D613.

Valk, P.J.M., Verhaak, R.G.W., Beijen, M.A., Erpelinck, C.A.J., Barjesteh van Waalwijk van Doorn-Khosrovani, S., Boer, J.M., Beverloo, H.B., Moorhouse, M.J., van der Spek, P.J., Löwenberg, B., et al. (2004). Prognostically useful gene-expression profiles in acute myeloid leukemia. N. Engl. J. Med. 350, 1617–1628.

Verhaak, R.G.W., Wouters, B.J., Erpelinck, C.A.J., Abbas, S., Beverloo, H.B., Lugthart, S., Löwenberg, B., Delwel, R., and Valk, P.J.M. (2009). Prediction of molecular subtypes in acute myeloid leukemia based on gene expression profiling. Haematologica 94, 131–134.

Wang, P., Ma, D., Wang, J., Fang, Q., Gao, R., Wu, W., Cao, L., Hu, X., Zhao, J., and Li, Y. (2016). INPP4B-mediated DNA repair pathway confers resistance to chemotherapy in acute myeloid leukemia. Tumour Biol. 37, 12513–12523.

Wouters, B.J., Löwenberg, B., Erpelinck-Verschueren, C.A.J., van Putten, W.L.J., Valk, P.J.M., and Delwel, R. (2009). Double CEBPA mutations, but not single CEBPA mutations, define a subgroup of acute myeloid leukemia with a distinctive gene expression profile that is uniquely associated with a favorable outcome. Blood 113, 3088–3091.

Yun, S., Vincelette, N.D., Yu, X., Watson, G.W., Fernandez, M.R., Yang, C., Hitosugi, T., Cheng, C.-H., Freischel, A.R., Zhang, L., et al. (2021). TFEB links MYC signaling to epigenetic control of myeloid differentiation and acute myeloid leukemia. Blood Cancer Discov. 2, 162–185.

Zhang, F., Zhu, J., Li, J., Zhu, F., and Zhang, P. (2017). IRF2-INPP4B axis participates in the development of acute myeloid leukemia by regulating cell growth and survival. Gene 627, 9–14.

Zhitomirsky, B., and Assaraf, Y.G. (2016). Lysosomes as mediators of drug resistance in cancer. Drug Resist Updat 24, 23–33.

Zuber, J., Radtke, I., Pardee, T.S., Zhao, Z., Rappaport, A.R., Luo, W., McCurrach, M.E., Yang, M.-M., Dolan, M.E., Kogan, S.C., et al. (2009). Mouse models of human AML accurately predict chemotherapy response. Genes Dev. 23, 877–889.

